# Glucose modulates transcription factor dimerization to enable tissue differentiation

**DOI:** 10.1101/2022.11.28.518222

**Authors:** Vanessa Lopez-Pajares, Aparna Bhaduri, Yang Zhao, Gayatri Gowrishankar, Laura Donohue, Margaret G. Guo, Zurab Siprashvili, Weili Miao, Duy T. Nguyen, Albert M. Li, Ronald L. Shanderson, Robin M. Meyers, Angela Guerrero, Andrew L. Ji, Omar S. Garcia, Shiying Tao, Lindsey M. Meservey, Xue Yang, Sanjiv S. Gambhir, Jiangbin Ye, Paul A. Khavari

## Abstract

Glucose is a universal energy currency for living organisms, however, its non-energetic functions in processes such as differentiation are undefined. In epidermis, differentiating cells exhibit dynamic changes in gene expression^1–4^ driven by specific transcription factors (TFs)^5–9^. The interplay between such TFs and biomolecules that also change in this process is not understood. Metabolomic analyses revealed that increased intracellular glucose accompanies differentiation of epidermal keratinocytes. This elevation also occurred in differentiating cells from other tissues and was verified in epidermal tissue engineered with glucose sensors, which detected a glucose gradient that peaked in the outermost differentiated layers. Free glucose accumulation, unaccompanied by its increased metabolism, was essential for epidermal differentiation and required GLUT1, GLUT3, and SGLT1 transporters. Glucose affinity chromatography and azido-glucose click chemistry uncovered glucose binding to diverse regulatory proteins, including the IRF6 TF, whose epidermal knockout confirmed its requirement in glucose-dependent differentiation. Direct glucose binding enabled IRF6 dimerization, DNA binding, genomic localization, and induction of IRF6 target genes, including essential pro-differentiation TFs *GRHL1, GRHL3, HOPX* and *PRDM1*. The IRF6^R84C^ mutant found in undifferentiated cancers was unable to bind glucose. These data identify a new role for glucose as a gradient morphogen that modulates protein multimerization in cellular differentiation.

---

Somatic tissue differentiation requires coordinated alterations in levels of the RNA transcripts, proteins, and other biomolecules necessary for tissue-specific function. This transformation has been characterized at the transcriptional level in a variety of cell types, including epidermal keratinocytes of skin^1–4^. Although specific TFs whose expression is induced during differentiation, such ZNF750, MAF, MAFB, GRHL3, HOPX, and PRDM1, are required for this process^1–3,5–9^, other TFs essential for differentiation are stably expressed. For example, the IRF6 TF, essential for epidermal development in mice^10,11^, is only modestly induced during differentiation^12^. Interplay between such TFs and potential molecular cues in this setting are incompletely understood. Transcriptional changes consistent with altered metabolism may be inferred in differentiating tissues but few studies have assessed how the metabolome itself changes^13,14^ in spite of the fact that metabolomics helps integrate transcriptomics and proteomics into a fuller picture of cellular function^15,16^. Glucose is an essential energy storage and transport molecule^17^ whose catabolism generates the ATP needed for diverse biologic functions. Major roles for glucose, independent of its energetic actions, are not established but indirect evidence suggests that glucose levels alter homeostatic processes. For example, the elevated glucose found in diabetes is associated with impaired fibroblast proliferation^18,19^ and high glucose itself was found to adversely impact stem cells^20^ and to promote osteogenic differentiation^21^. Additionally, elevated glucose is associated with impaired skeletal muscle progenitor proliferation^22^. Potential non-energetic mechanisms whereby glucose levels might control differentiation in self-renewing tissues, however, are not well characterized.

An unbiased analysis was performed of intracellular analytes during human epidermal keratinocyte differentiation. 5 types of mass spectrometry (**MS**), including UPLC-MS, positive and negative ionization LC-MS, LC-polar-MS, and GC-MS, capable of detecting >14,000 analytes in major metabolite classes were used. Analyses were performed on 5 independent biological replicates each in a) proliferating undifferentiated progenitor-containing populations in vitro as well as keratinocytes differentiated at confluence with calcium b) for 3 days to represent early differentiation, and c) for 6 days to represent late differentiation^23^. Of 614 identified metabolites (**Fig. 1a, Table S1**), 193 significantly changed during this process. Among increased analytes were those previously associated with terminal differentiation in skin, including γ-glutamyl amino acid substrates of transglutaminase important in cutaneous barrier formation^24^. Also elevated were fatty acids, which promote skin differentiation by activating PPARα and by contributing to cutaneous barrier function^25^ (**Extended Data Fig. 1a-b**). The top increased analyte during epidermal differentiation was trans-urocanate (**Extended Data Fig. 1c**), which comprises 0.5% of the dry weight of the stratum corneum, where it may provide photoprotection^26^. Detection of these known differentiation-induced analytes supports the biologic relevance of this dataset.

**Fig 1.**
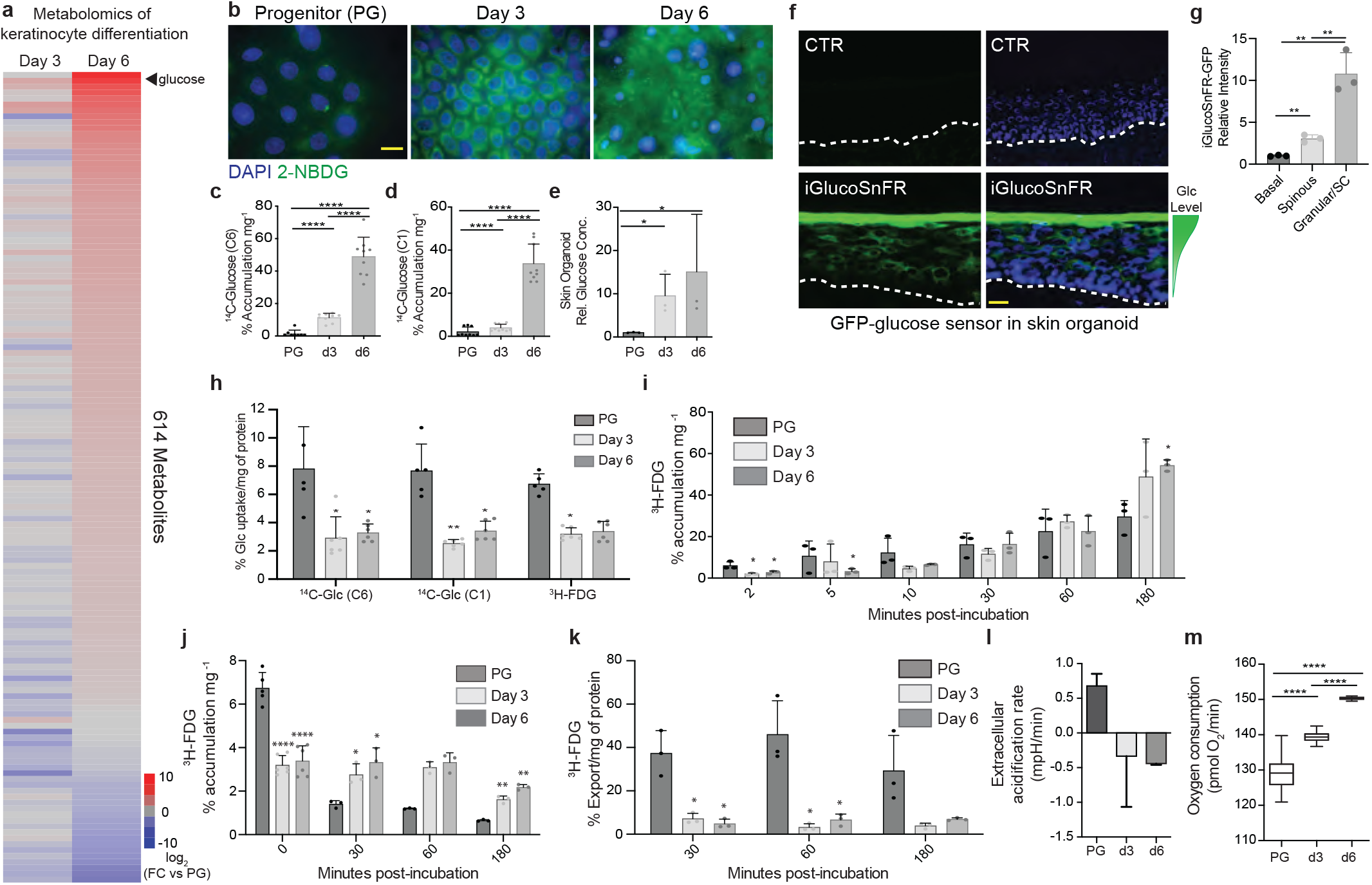
Intracellular glucose accumulates in keratinocyte differentiation. **(a)** Heatmap representing the 614 metabolites identified in metabolomics, shown as a fold change at early (d3) and late (d6) stages of differentiation compared to the undifferentiated state. The average of 5 biological replicates is shown. **(b)** 2-NBDG, a fluorescent glucose analog, accumulates in differentiated keratinocytes; 2-NBDG [green], Hoechst nuclear stain [blue]. Progenitor (PG) represents undifferentiated cells vs d3 and d6 of calcium-induced differentiation in vitro (representative image of 3 independent samples, scale bar=10μm). Quantitation of radiolabeled glucose Glucose-C14 (C6) **(c)** or Glucose-C14 (C1) **(d)** accumulation during differentiation in culture, incubated for 7 d, normalized to mg of protein in each sample (n=3 biological replicates and 3 technical replicates per experiment). **(e)** Quantitation of glucose concentrations in regenerated differentiating organoid epidermal tissue; whole epidermis undergoing stratification through d6 analyzed (n=3 biological replicates). **(f)** Epidermal organoids expressing vector control or the GFP glucose sensor, iGlucoSnFR, for 7d. The GFP signal [green] indicates detection of free glucose in the tissue, with nuclear DAPI stain [blue] (representative image of 3 biological replicates, scale bar=20μm). **(g)** Quantitation of GFP signal in the basal layer, spinous layer or stratum corneum of the tissue was measured (n=3 biological replicates). **(h)** Radiolabeled glucose uptake in PG and differentiating (d3, d6) keratinocytes after 5 minutes of incubation with the tracer normalized to the amount of protein in each sample (n=3 biological replicates, 3 technical replicates per experiment). **(i)** ^3^H-FDG accumulation in PG and differentiated (d3, d6) keratinocytes over a time course of 2 minutes to 3 hours (n=3 biological replicates, 3 technical replicates per experiment). **(j)** Percent accumulation of glucose assayed at 30 min and 1 h after PG and differentiated (d3, d6) cells were incubated with ^3^H-FDG for 5 minutes (n=3 biological replicates, 3 technical replicates per experiment). **(k)** Percent exported ^3^H-FDG assayed at 30 min, 1 h and 3 h after cells were incubated with ^3^H-FDG for 5 min. Export was calculated by measuring media for the tracer normalized to the amount of protein in each sample (n=3 biological replicates, 3 technical replicates per experiment). **(l)** Glycolysis rates were measured in PG or differentiation (d3, d6) via the extracellular acidification rate (ECAR) (n=3 biological replicates). **(m)** Oxygen consumption was measured in PG and differentiated (d3, d6) keratinocytes (n=3 biological replicates). Statistical significance was determined using unpaired two-tailed Student’s *t*-test. Error bars represent S.D., *p<0.05, **p<0.01, ***p<0.005, ****p<0.001.

Unexpectedly, the second most increased analyte in keratinocyte differentiation was free glucose (**Extended Data Fig. 1d**). This finding was validated using four orthogonal methods. First, the fluorescent glucose analog, 2-[N-(7-nitrobenz-2-oxa-1,3-diazol-4-yl) amino]-2-deoxy-D-glucose (**2-NBDG**), which undergoes uptake at rates similar to unlabeled D-glucose^27^, was incubated with differentiating epidermal cells in vitro. 2-NBDG loses fluorescence upon phosphorylation, allowing detection of free, non-metabolized glucose^28^. Epidermal cells increased intracellular 2-NBDG during differentiation, consistent with free glucose accumulation (**Fig. 1b, Extended Data Fig. 1e**). Second, ^14^C-radiolabeled glucose tracers were used. Glucose can be metabolized by multiple catabolic pathways, including the glycolysis and pentose phosphate pathways, however, glucose accumulation was observed when using both ^14^C-glucose (C6), which is metabolized via glycolysis (**Fig. 1c**), and ^14^C-glucose (C1), which is metabolized via the pentose phosphate pathway (**Fig. 1d**). Third, ^3^H-Fludeoxyglucose (^3^H-FDG), which can be phosphorylated but not metabolized^29^, showed the same trend (**Extended Data Fig. 1f**). Finally, the iGlucoSnFR glucose sensor, which fluoresces in the presence of free glucose,^30–32^ was used. In live-cell imaging during keratinocyte differentiation in vitro, iGlucoSnFR detected marked increases in glucose, which were impaired in low glucose media (**Extended Data Fig. 1g-h**). Nuclear iGlucoSnFR detected elevated glucose in differentiating keratinocyte nuclei as well (**Extended Data Fig. 1i**). To examine glucose accumulation in 3-dimensional (**3D**) tissue, iGlucoSnFR was expressed in human skin organoids using an approach that recapitulates the 3D architecture and gene expression of human skin tissue^2,3,23,33^. iGlucoSnFR detected increased glucose in outer differentiating layers of epidermal tissue, consistent with glucose quantitation in differentiating organoids (**Fig. 1e-g, Extended Data Fig. 1j-k**). Glucose also accumulated within differentiating adipocytes, myoblasts and osteoblasts in vitro (**Extended Data Fig. 1l-p**) indicating that, although the magnitude of glucose increases may vary based on the cellular context and assay used, elevation of intracellular glucose occurs during the differentiation of diverse somatic cell types.

The basis for differentiation-associated glucose accumulation was next explored, beginning with analysis of glucose uptake and export from differentiating keratinocytes. Radiolabeled glucose displayed decreased uptake as cells differentiated (**Fig. 1h**). Consistent with this, progenitor cells increased intracellular ^3^H-FDG within 5 minutes, however, paradoxically, by 3 hours differentiating cells contained twice as much ^3^H-FDG as progenitors (**Fig. 1i**). Thus, despite decreased uptake, differentiating cells accumulate intracellular glucose by exporting less glucose than undifferentiated cells. Indeed, an ^3^H-FDG pulse-chase experiment demonstrated that intracellular ^3^H-FDG decreased dramatically over time in progenitor cells but not differentiating cells (**Fig. 1j**). Additionally, more ^3^H-FDG was detected in media from progenitors post wash-out compared to the differentiated cells (**Fig. 1k**), indicating that differentiating keratinocytes accumulate glucose in concert with decreased glucose export.

A possible reason for glucose accumulation in differentiation might be for use in glycolysis or other metabolic processes. Extracellular acidification rates, a surrogate measure for glycolysis, however, indicated that glycolytic rates decreased during differentiation (**Fig. 1l**). This was accompanied by a significant increase in oxygen consumption, consistent with a shift away from glycolysis towards oxidative phosphorylation (**Fig. 1m**) characteristic of differentiating cells^34^. Moreover, 18 major pathway metabolite intermediates in all three major glucose catabolic pathways, namely the glycolysis, pentose phosphate, and hexosamine pathways, did not accumulate during differentiation (**Extended Data Fig. 2a-b)** nor did the hexosamine pathway product, β-linked *N*-acetylglucosamine (**O-GlcNAc**) (**Extended Data Fig. 2d-e**).

To gain further insight into cellular energetics relevant to glucose during differentiation, metabolic flux analysis was performed on progenitor, day 3 and day 6 differentiating keratinocytes. Consistent with metabolomic data, [U-^13^C]-glucose incorporation into glycolysis intermediates decreased during differentiation (**Extended Data Fig. 2f-g**) and lactate production, was not significantly increased in late differentiation (**Extended Data Fig. 2h-i**). Notably, a significant increase in [U-^13^C]-glucose conversion to α-ketoglutarate was observed during differentiation, a finding that correlates with the observed increase in oxidative phosphorylation and is consistent with a metabolic shift from glycolysis to glucose oxidation characteristic of differentiated cells^34,35^.

The contribution of fatty acids to cellular energetics during differentiation was evaluated by interrogating metabolites related to fatty acid metabolism in the metabolomics data. A modest increase in various fatty acid metabolites was found, including a significant increase in acetylcarnitine, a metabolite required for fatty acid β-oxidation (**Extended Data Fig. 2k)**. Consistent with these findings and enrichment for fatty acids in the metabolomics data in early differentiation, protein expression of lipid metabolism regulators, such as ACSL1 and ATP-citrate lysase, was found elevated in progenitor and d3 differentiation (**Extended Data Fig. 2l)**, suggesting an early contribution of lipid metabolism towards differentiation. Together, these findings demonstrate that intracellular free glucose accumulates without evidence for increased glucose catabolism.

To assess functional impacts of glucose elevation, keratinocytes were incubated in either standard media extracellular glucose concentrations of 4.5g/L (25mM) or in a reduced glucose media of 0.5 g/L (2.8mM) that still supports normal cell viability. Keratinocytes proliferated normally at both concentrations, however, cell growth ceased at 0 g/L glucose (**Extended Data Fig. 3a**), consistent with a glucose requirement for cellular energetics^36^. Compared to standard glucose media, glucose restriction decreased intracellular keratinocyte glucose concentrations and suppressed epidermal differentiation gene expression (**Extended Data Fig. 3b-c**). Similarly, human skin organoids grown in low glucose media displayed impaired differentiation (**Fig. 2a-b**). RNA-sequencing of replicate epidermal organoid tissues (n=4/group) cultured in standard or low glucose media identified 3,208 differentially expressed genes. Foremost among genes downregulated in low glucose were those implicated in epidermal differentiation (**Fig. 2c-d, Table S2**). Significant overlap occurred between glucose-modulated genes and previously identified^2,3^ epidermal differentiation gene signatures (**Fig. 2e**). The 3-O-methyl-glucose (**3MG**) glucose analog, which is metabolically stable in cells and tissues^37^, rescued differentiation of cells grown in low glucose media (**Fig. 2f**). These findings indicate that glucose levels beyond those necessary for normal cell viability and proliferation are essential for keratinocyte differentiation and suggest that free glucose itself, as opposed to its catabolism, plays a role in this process.

**Fig 2.**
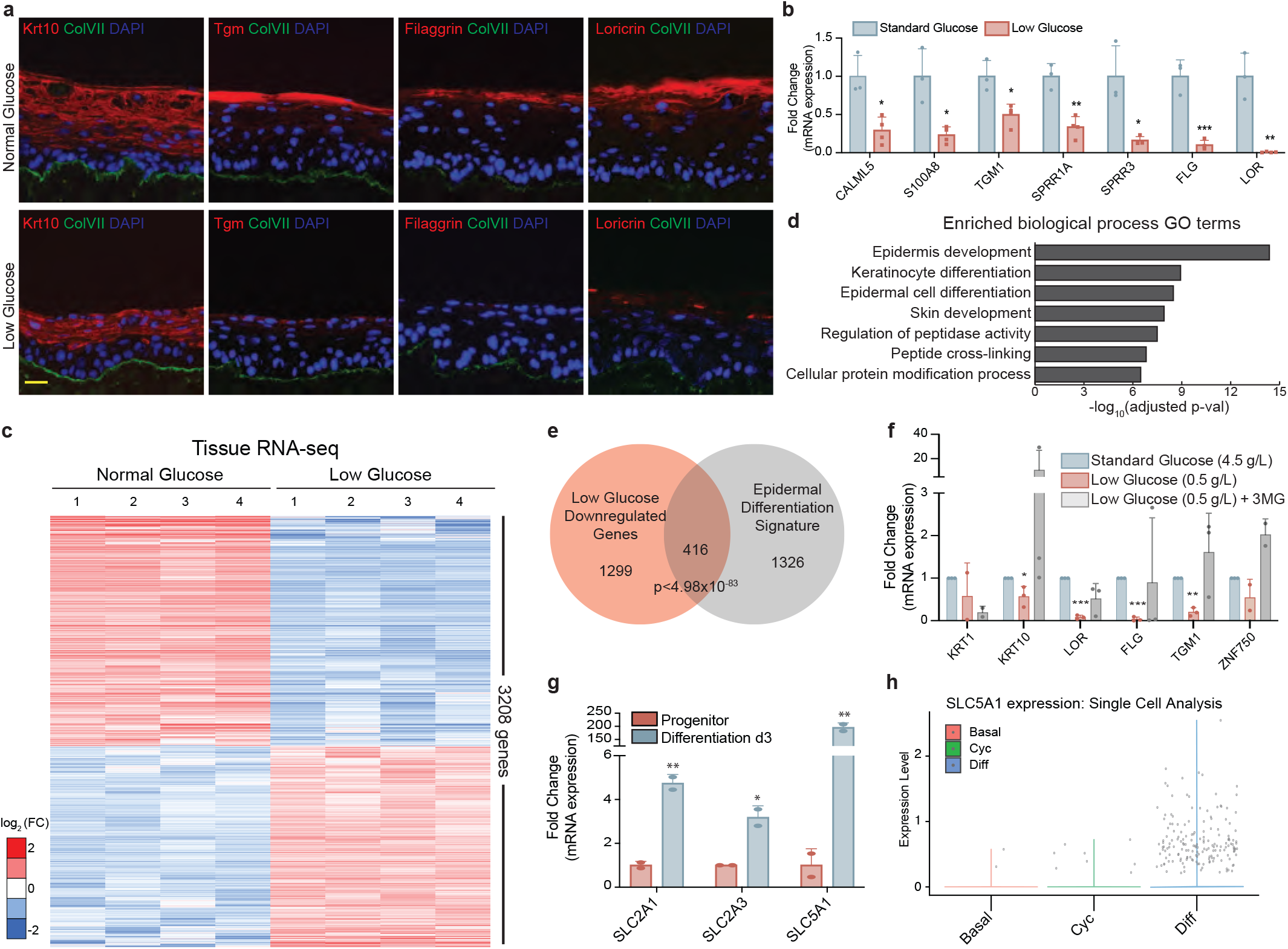
Glucose is required for epidermal differentiation. **(a)** Epidermal organoids grown in standard (4.5g/L; 25mM) or low (0.5g/L; 2.8mM) glucose media; differentiation markers [red], nuclei [blue], collagen VII basement membrane staining [green] (representative images shown, n=4 biological replicates,). Normalized differentiation gene mRNA expression from epidermal organoids in (A) (n=4 biological replicate organoids/condition). **(c)** Heatmap of 3208 differentially expressed genes (PAS-seq) of epidermal organoids in (A) (n=4 biological replicates, FDR=0.05, fold change>1.5). **(d)** Enriched Gene Ontology terms of the downregulated genes in low glucose media relative to standard glucose media. **(e)** Overlap of genes downregulated in low glucose media with the epidermal differentiation signature genes from Rubin et al. and Lopez-Pajares et al. **(f)** Normalized gene expression of differentiated (d3) keratinocytes grown in standard glucose, low glucose, or low glucose supplemented with the 3-O-methyl-glucose analog (**3MG**) to rescue glucose levels (n=3 biological replicates). **(g)** mRNA expression of glucose transporters in progenitor and d3 differentiated keratinocytes measured by qPCR (n=2 biological replicates). **(h)** mRNA expression levels of *SLC5A1* (SGLT1) in basal, cycling or differentiated cell populations from single-cell analysis of normal skin. Statistical significance was determined using unpaired two-tailed Student *t*-test. Error bars represent S.D. (b,f) or S.E.M. (g), *p<0.05, **p<0.01, ***p<0.005.

To further assess a requirement for accumulated free glucose in differentiation, intracellular glucose was reduced by expressing glucose metabolizing enzymes that lower intracellular glucose concentrations. Epidermal organoid tissue with enforced expression of Hexokinase 1 and 2 (**HK1/2**) displayed decreased intracellular glucose and impaired differentiation (**Extended Data Fig. 3d-h**). To verify this change was driven by HK1/2 enzymatic activity, the 2-deoxyglucose (2DG) HK1/2 competitive binding inhibitor was used^38^. 2-DG partially restored intracellular glucose levels and rescued differentiation markers decreased by enforced HK1/2 expression (**Extended Data Fig. 3i-j**). Enforced expression of G6PD, the glucose metabolic enzyme responsible for the first steps of the pentose phosphate pathway, likewise lowered intracellular glucose and impaired tissue differentiation (**Extended Data Fig. 3k-l**). This phenotype was also successfully rescued by a non-competitive G6PD inhibitor^39^ that restored intracellular glucose levels and differentiation gene expression (**Extended Data Fig. 3m-n**). These data are consistent with a model in which accumulated intracellular glucose itself is required for epidermal differentiation.

The contribution of specific glucose transporters^40^ to glucose accumulation during differentiation were next examined. The primary glucose transporters, GLUT1 (*SCL2A1*) and to a lesser extent GLUT3 (*SLC2A3*), were modestly increased in differentiated keratinocytes, whereas the sodium-coupled glucose co-transporter, SGLT1 (*SLC5A1*), increased >50-fold (**Fig. 2g, Extended Data Fig. 4a-c**). In single-cell analysis of samples of normal adult human skin^41^, *SLC5A1* was expressed almost exclusively in differentiating keratinocytes (**Fig. 2h, Extended Data Fig. 4d-e**). Several pro-differentiation transcription factors, including p63, ZNF750, KLF4, CEBPα/β/δ and IRF6, were found to regulate gene expression of these transporters (**Extended Data Fig. 4f, Table S3**). Furthermore, glucose transporter protein detection correlated with gene expression in progenitor and differentiated keratinocytes; GLUT1 and GLUT3 were largely unchanged, whereas SGLT1 expression increased and was re-localized diffusely throughout the cell (**Extended Data Fig. 4g**). To define contributions of these glucose transporters to the accumulation of glucose during differentiation, independent CRISPR-Cas9-mediated knockouts of *SLC2A1, SLC2A3* and *SLC5A1* genes was performed. Each showed impaired differentiation (**Extended Data Fig. 4h-l**). Consistent with this, differentiating keratinocytes treated with pharmacological inhibitors WZB117, targeting the GLUT family of transporters, or phlorizin, targeting the SGLT family of transporters resulted in decreased intracellular glucose and suppressed differentiation gene expression (**Extended Data Fig. 4m-n**). These data support non-redundant roles for multiple glucose-transporters, including the differentiation-upregulated SGLT1 symporter, in mediating the accumulation of intracellular glucose in differentiating keratinocytes.

Glucose binds specific proteins, such as HK1/2, raising the possibility that elevated free glucose might bind proteins important for differentiation. To search for these, extracts from differentiating keratinocytes were passed over a glucose polymer column then eluted with either free glucose or galactose monosaccharide control and subjected to LC-MS/MS. The 497 proteins enriched in the glucose elution (**Table S4**) were concentrated in biological processes related to gene expression, translation, and protein metabolism (**Extended Data Fig. 5a-b**). 18 glucose-associated proteins were identified with ≥2-fold mRNA induction during differentiation (**Fig. 3a**), including the IRF6 TF.

**Fig 3.**
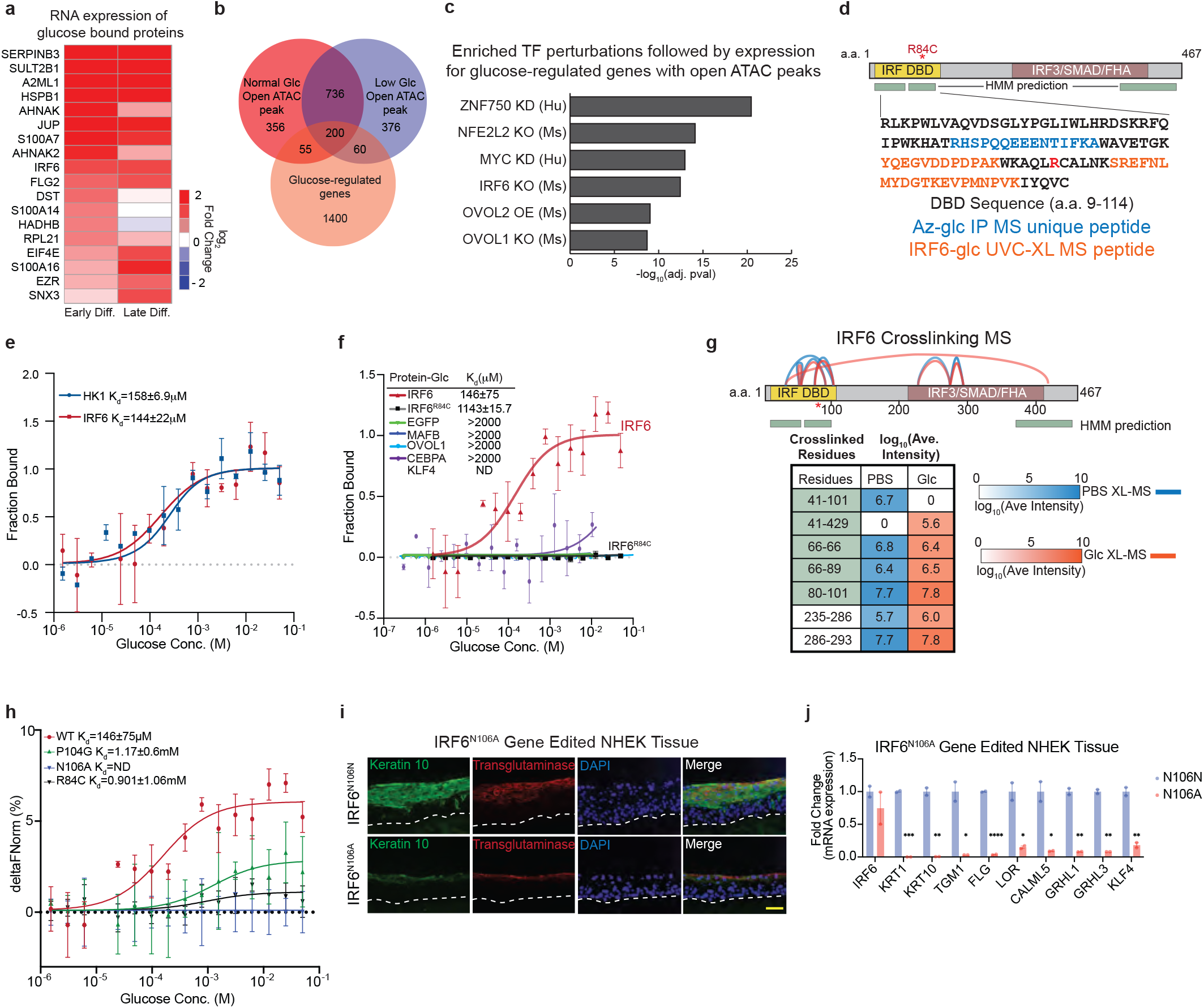
Identification of IRF6 as a glucose binding protein. **(a)** Gene expression of differentiation-induced glucose-binding proteins identified via affinity chromatography followed by LC-MS/MS. **(b)** Overlap of glucose-regulated genes and ATAC-seq connected genes with increased accessibility (open peak) in d3 differentiated keratinocytes grown in standard (n=1347) or low glucose (n=1372) relative to progenitor keratinocytes. **(c)** Enriched TF target gene sets after perturbation in glucose-regulated genes (maayanlab.cloud/Enrichr). **(d)** Schematic of IRF6 protein domains indicating DNA binding domain (DBD) and IRF3 homology domain. The highlighted sequence in the DNA binding domain represents the unique peptide identified in azido-glucose pulldown followed by MS and glucose-binding peptides identified by UVC crosslinking MS. In red is the *R84C mutation, which is found in IRF6-driven developmental disorders and in cancer. Green bars represent HMM predicted glucose binding regions. **(e)** MST analysis was used to determine the affinity of glucose toward HK1 (K_d_=158+/−6.9μM) or IRF6 (K_d_=144+/−22μM) (n=4). **(f)** MST analysis to determine the affinity of glucoses toward an EGFP control or transcription factors IRF6 (K_d_=146+/−0.6μM), IRF6^R84C^ mutant (K_d_=1143+/−1.57μM), CEBPA, OVOL1, MAFB, and KLF4 (K_d_>2mM or not detected) (n=3). **(g)** Chemical crosslinking followed by MS identified crosslinked residues and shown are the average intensity values for all the crosslinks observed during IRF6 incubation with PBS (blue) or 350μM glucose (orange). *Denotes location of R84C mutation. Green bars/shading indicate HMM predicted glucose binding regions in IRF6. **(h)** MST analysis was used to determine the affinity of glucose toward IRF6^P104G^ (K_d_=1.17+/−0.6mM) or IRF6^N106A^ (K_d_ not detected) compared to wild-type and IRF6^R84C^ (n=4). **(i)** Epidermal organoids generated from IRF6^N106N^ and IRF6^N106A^ gene edited keratinocytes; differentiation markers keratin 10 [green], transglutaminase [red], and nuclear DAPI stain [blue] (representative image of 2 biological replicates, scale bar=50μm). **(j)** qPCR gene expression analysis of tissue in (i) (n=2 biological replicates, representative data for 2 technical replicates shown).. Statistical significance determined using unpaired two-tailed Student’s *t-*test. Error bars represent S.D. (e,f,h) or S.E.M. (j), *p<0.05, **p<0.01, ***p<0.005.

Epidermal differentiation is accompanied by dynamic changes in genome accessibility at thousands of regulatory elements across the genome^23^. To explore glucose impacts on this, ATAC-seq was performed on differentiating keratinocytes grown in standard versus low glucose media. Glucose restriction decreased chromatin accessibility at putative regulatory regions of 406 genes, including gene promoters and H3K27ac-marked distal promoter-interacting regions (**PIRs**) assigned by H3K27ac HiChIP data obtained in differentiating keratinocytes^23^; glucose restriction also increased accessibility at 436 genes (**Fig. 3b, Table S5**). To identify glucose-regulated genes, genes with increased accessibility in standard glucose media compared to glucose restriction were intersected with the 1,715 genes whose RNA transcripts were downregulated with low glucose (**Fig. 3b**). The resulting 55 candidate direct glucose target genes were enriched for biologic actions in epidermal differentiation and skin development (**Extended Data Fig. 5c**). These genes and their corresponding regulatory regions were then used to nominate candidate TFs involved in glucose-mediated gene regulation. Among identified TF candidates were those known to be essential for normal epidermal differentiation (**Fig. 3c**), including ZNF750, OVOL1, and IRF6.

Several studies have shown that IRF6 is essential for epidermal development in mice^10,11,42^. The impact of CRISPR-Cas9 mediated *IRF6* deletion was therefore assessed in human epidermal organoid tissues. Four different sgRNAs targeted *IRF6* in primary keratinocytes used to generate epidermal organoids. Consistent with its mouse phenotype, *IRF6* targeting impaired epidermal differentiation (**Extended Data Fig. 5d-g**). RNA-sequencing of biological replicates of IRF6 depleted organoid tissue identified 1,191 downregulated and 720 upregulated genes (**Extended Data Fig. 5h, Table S6**). Downregulated genes were enriched for skin development, epidermal differentiation and sphingolipid metabolism, similar to the phenotypes observed in *Irf6* knockout mice^10,42^ (**Extended Data Fig. 5i**). Significant overlap was found between IRF6-regulated genes, epidermal differentiation signatures, and the glucose-dependent differentiation gene set (**Extended Data Fig. 5j**), consistent with the analysis above. These findings verify conservation of IRF6 function in epidermal differentiation cross-species and raise the possibility that IRF6 mediates a portion of the glucose-enabled differentiation program.

Having identified IRF6 as a putative glucose binding protein, click-chemistry with an azido-conjugated glucose analog, 2-azido-2-deoxy-glucose, was next performed as an orthogonal method to glucose polymer affinity. LC-MS/MS identified a unique peptide enriched in the azido-glucose pull-down in the IRF6 DNA binding domain (**Fig. 3d, Table S7**), an amino-terminal portion of IRF6 that Hidden Markov Modeling (**HMM**) predicted as a glucose-binding region (**Fig. 3d**, green bars). Microscale thermophoresis (**MST**) showed that purified recombinant IRF6 protein binds glucose with a dissociation constant of 144±22μM, similar to HK1 (*K*_*d*_=158±6.9μM) (**Fig. 3e**). IRF6 also bound the glucose analog 3MG, shown capable of rescuing differentiation above, at comparable affinity (**Extended Data Fig. 6a**). The glucose-TF interaction was selective for IRF6, because recombinant pro-differentiation TFs CEBPA, MAFB, OVOL1 and KLF4 failed to bind glucose (**Fig. 3f**). The IRF6^R84C^ point mutation, which is a recurrent somatic alteration in undifferentiated cancers and whose germline presence causes Van der Woude and popliteal pterygium syndromes^43,44^ reduced IRF6 binding to glucose by 100-fold (**Fig. 3f**).

The impact of glucose binding on IRF6 conformation was next assessed by chemical crosslinking-MS in the presence of PBS or 350μM glucose; the latter is the physiologic intracellular concentration in differentiating keratinocytes (**Extended Data Fig. 6b**). Several crosslinks were identified within the IRF6 DNA binding domain and within the IRF3/SMAD/FHA domain in both conditions, however, a specific crosslink between lysine 41 and lysine 101 within the DNA binding domain present in PBS only was lost with glucose and instead lysine 41 was found crosslinked to lysine 429, indicating that glucose modulates the conformation of IRF6 (**Fig. 3g, Table S7**). Interestingly, a crosslink at lysine 66 was identified between two different IRF6 protein molecules, indicating that glucose may facilitate IRF6 dimer formation in vitro.

To identify the direct glucose binding residues of IRF6, UVC crosslinking-MS was performed with recombinant IRF6 protein, with or without incubation with glucose. Peptides mapping to the DNA binding domain and C-terminus of IRF6 were identified as putative glucose binding sites and glucose was found crosslinked at residues P104 and N106, residing within the DNA binding domain (**Fig. 3d, Extended Data Fig. 6c-d)**. In agreement with this, molecular docking studies^45,46^ with glucose and an AlphaFold predicted IRF6 structure^47^ predicted binding free energy of −5.1 kcal/mol, placing glucose in 3D space adjacent to the IRF6 peptide (a.a. 45-59) identified via azido-glucose pulldown, the direct glucose-binding residues P014 and N106, as well as R84 (**Extended Data Fig. 6e)**. These data indicate that glucose directly binds IRF6 in a fashion dependent intactness of specific amino acids within the IRF6 amino terminal region.

The contribution of glucose-binding residues to IRF6 function was next explored. First, MST analysis was used to demonstrate that mutation of IRF6 P104G resulted in decreased affinity towards glucose, while mutation N106A abolished the interaction, similar to R84C (**Fig. 3h)**. These mutants were further characterized by an electrophoretic mobility shift assay (EMSA) to assess their DNA binding capacity. The IRF6^P104G^ mutant lost the ability to bind DNA, while this function remained intact for IRF6^N106A^ (**Extended Data Fig. 6f**). Enforced expression of these mutants, and a P104G/N106A double mutant, in the context of *IRF6* depletion failed to rescue differentiation gene expression to the extent of wild-type IRF6 enforced expression (**Extended Data Fig. 6g-h**). CRISPR-Cas9/AAV mediated homology-directed recombination was next employed to perform gene editing of the endogenous *IRF6* locus in normal human epidermal keratinocytes. Gene editing of *IRF6* to mutate R84C, P104G and N106A was effected at >90% efficiency in pooled normal diploid human keratinocytes (**Extended Data Fig. 6i**). These cells were used to generate epidermal organoids, which revealed a requirement for glucose binding to fully induce differentiation gene expression (**Fig. 3i-j, Extended Data Fig.6 j-m**). Notably, the IRF6^N106A^ mutant, which retains DNA binding capacity but cannot bind glucose, displayed impaired differentiation gene expression comparable to the DNA-binding deficient mutants, establishing a critical role for glucose in IRF6-dependent gene expression.

The finding that a subset of glucose-regulated genes are IRF6-dependent focused efforts on the mechanism by which glucose impacts IRF6 gene regulation. Because TFs such as IRF6 exert their regulatory impacts by associating with DNA, the capacity of glucose to effect IRF6 DNA binding was first examined. IRF6 was incubated with DNA oligonucleotides derived from endogenous genomic sequence that contained an IRF6 binding motif in regulatory DNA at the *DMKN* differentiation gene locus, a known IRF6 differentiation target, in the presence or absence of the 350μM glucose concentration present intracellularly in differentiated keratinocytes. Glucose enhanced IRF6 binding to its cognate DNA binding sequence by 151-fold (*K*_*d*_ = 1.21±0.86μM in PBS to *K*_*d*_ = 0.008±0.00714μM in glucose). In contrast, glucose had minimal impact on the low affinity association observed for IRF6 to a mutant *DMKN* oligo lacking an intact IRF6 motif (**Fig. 4a**), indicating that glucose enhances IRF6 binding to its cognate DNA sequence motifs but not to DNA in general. A similar effect was observed for IRF6 binding to a native IRF6 binding site in the *SFN* differentiation gene locus, where a 45-fold increase in IRF6 DNA binding affinity with glucose was quantified (*K*_*d*_ = 16.4±5.8μM in PBS to *K*_*d*_ = 0.357±0.215μM in glucose) (**Extended Data Fig. 7a-b**). Glucose had no impact on DNA binding by MAFB to a native MAFB binding site at the *GRHL3* differentiation gene locus (**Extended Data Fig. 7c**), further indicating that glucose lacks non-specific global effects on TF-DNA associations. EMSA and DNA ELISA of recombinant IRF6 incubated with 350μM glucose, but not the 100μM glucose concentration present in progenitor keratinocytes, demonstrated enhanced DNA binding activity (**Extended Data Fig. 7d-e**). These data demonstrate that glucose increases the association of IRF6 with DNA.

**Fig 4.**
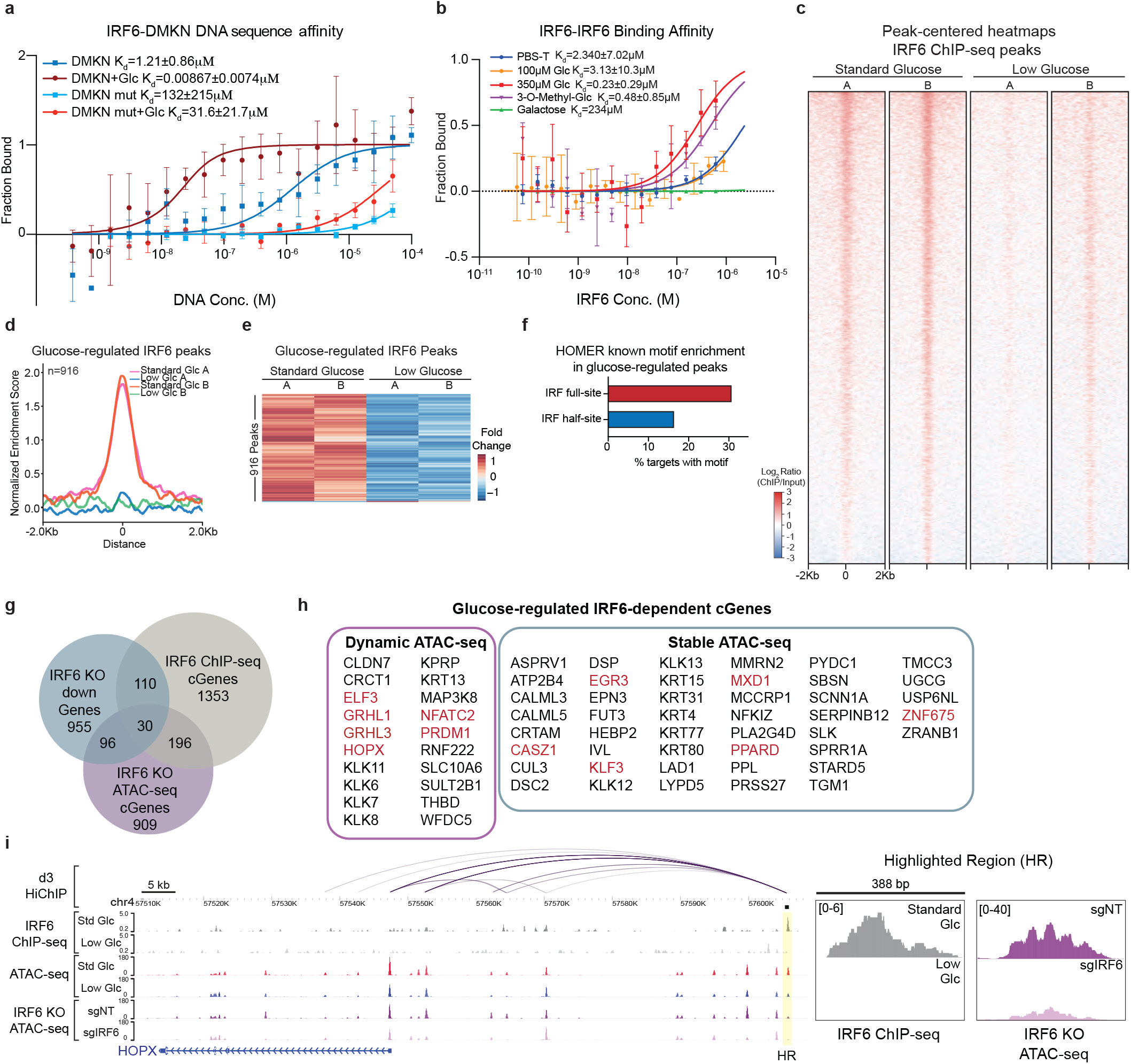
Glucose enhances IRF6 DNA binding and modulates gene regulation. **(a)** Affinity of IRF6 towards cognate DNA binding sequence in the *DMKN* locus measured by MST. Measurements were conducted after incubating IRF6 with a wild-type *DMKN* sequence oligo in PBS (K_d_=1.21±0.86μM) or in 350μM glucose (K_d_=0.00867±0.00714μM) or an IRF6 binding site mutated *DMKN* oligo in PBS (K_d_=137±215μM) or with 350μM glucose (K_d_=31.6±21.7μM) (n=4). **(b)** MST analysis of IRF6 affinity toward another IRF6 molecule to assess oligomerization in the presence of PBS (K_d_=2.34±7.02μM), 100μM glucose (K_d_=3.13±10.3μM), 350μM glucose (K_d_=0.23±0.29μM), 350μM 3-O-methyl-glucose (K_d_=0.48±0.85μM) or 350μM galactose (K_d_=234μM) (n=4). **(c)** Peak-centered heatmap of IRF6 ChIP-seq peaks in standard (11649 peaks) or low (7469 peaks) glucose media d3 differentiated keratinocytes (n=2 biological replicates/condition). **(d)** Normalized enrichment score for 916 glucose-regulated IRF6 ChIP-seq peaks in standard or low glucose (n=2 biological replicates/condition). **(e)** Heatmap of 916 differential IRF6 ChIP-seq peak in d3 differentiated keratinocytes in standard glucose compared to low glucose media. **(f)** HOMER known motif analysis showing percentage of targets with IRF4 motif, representing the IRF half-site, or IRF full-site, consisting of motifs matching IRF2, IRF3, IRF8 or ISRE, in glucose-regulated IRF6 ChIP-seq peaks. **(g)** Identification of putative IRF6 direct targets by overlap of IRF6 cGenes in differentiated keratinocytes (d3) assigned to IRF6 ChIP-seq peaks or IRF6 KO ATAC-seq cGenes and IRF6 KO tissue downregulated genes. **(h)** Table of glucose-regulated IRF6 genes with dynamic or stable ATAC-seq peaks in IRF6 KO differentiated keratinocytes (d3); TFs are highlighted in red. **(i)** Genome browser tracks of d3 HiChIP, IRF6 ChIP-seq peaks and ATAC-seq peaks in differentiated keratinocytes (d3) in standard or low glucose media, and ATAC-seq peaks in control NT or IRF6 KO keratinocytes (d3) for the *HOPX* locus. All error bars represent S.D.

Efficient DNA binding and gene regulation by IRF family TFs can depend on IRF protein dimer formation^48^, therefore we next tested the ability of glucose to mediate IRF6 homodimerization. Consistent with cross-linking MS data above, 350μM glucose increased IRF6 dimerization, as did 3MG, but neither galactose nor 100 μM glucose, boosted IRF6 dimerization (**Fig. 4b**). These glucose impacts were orthogonally verified using co-immunoprecipitation with V5 and HA epitope tagged IRF6 proteins expressed in differentiating keratinocytes grown in standard or low glucose media where glucose also enhanced IRF6 dimer formation (**Extended Data Fig. 7f-g**); IRF6 dimerization was mildly observed in the cytoplasmic, but not in the nuclear fraction of progenitor keratinocytes (**Extended Data Fig. 7h)**. The IRF6^S413/424A^ dimerization-deficient mutant, modestly diminished dimerization in both conditions (**Extended Data Fig. 7i**). These data indicate that glucose facilitates IRF6 dimer formation, an important process in IRF family TF binding to DNA.

The impact of glucose on IRF6 nuclear activity was next assessed. Consistent with glucose enhancement of IRF6 DNA binding, an IRF6 reporter displayed greater luciferase activity in differentiating keratinocytes grown in standard versus low glucose media in vitro (**Extended Data Fig. 7j**). No substantial differences in nuclear IRF6 were seen between standard versus low glucose media, suggesting that glucose does not alter IRF6 subcellular localization (**Extended Data Fig. 7k-l**). To examine glucose impacts on IRF6 genomic localization, biologic replicate IRF6 ChIP-seq were next performed in differentiating keratinocytes grown in standard or low glucose media. 11,649 IRF6 peaks were identified in standard glucose media, with a substantially reduced number of 7,469 peaks identified with glucose restriction; stronger normalized peak enrichments were seen with glucose (**Fig. 4c, Extended Data Fig. 8a, Table S8**). Potential sites of active gene regulation found in H3K27Ac HiChIP data from differentiating human keratinocytes^23^ that were in physical contact with IRF6-bound regions identified by ChIP-seq nominated putative IRF6-connected target genes (cGenes). This identified 3,858 IRF6 cGenes in standard glucose media and 1,988 cGenes with glucose restriction; 1,807 of these cGenes were shared between the two conditions. Thus, substantially more candidate IRF6-regulated genes were identified with standard glucose (**Extended Data Fig. 8b**). *De novo* motif analysis identified IRF6 motif enrichment in IRF6 peaks shared between the standard and low glucose conditions (**Extended Data Fig. 8c**). Differential analysis to define the IRF6 peaks controlled by glucose identified 916 peaks, of which only 8 were downregulated in standard glucose (**Fig. 4d-e, Table S8**), consistent with the observation in vitro that glucose enhances the binding of recombinant IRF6 protein to its cognate DNA sequences. The 908 glucose upregulated peaks were enriched for epidermal differentiation related GO terms (**Extended Data Fig. 8d**), further supporting a role for glucose in promoting IRF6-mediated pro-differentiation gene expression. The top three enriched motifs in HOMER de novo motif analysis of glucose-regulated peaks included BATF, KLF4 and IRF4 (**Extended Data Fig. 8e**). Glucose-regulated IRF6 binding peaks were more abundant in promoter features within 1Kb of the transcription start site, compared to low glucose, which concentrated in intergenic or intronic regions (**Extended Data Fig. 8f-g**), suggesting that glucose redirects IRF6 to potentially activating gene regulatory elements. Low glucose IRF6 peaks lacked statistical enrichment for IRF motifs, in contrast to glucose-regulated peaks, which showed strong IRF motif enrichment. The latter showed enrichment for an IRF4 motif, which, like the IRF6 motif, contains a single core GAAA sequence (half-site), with a higher overall percentage of peaks that contained a motif resembling IRF family members that bind as dimers (full-site IRF2, IRF3, IRF8, ISRE motifs) (**Fig. 4f**). These results support in vitro findings showing that glucose promotes IRF6 dimerization and DNA binding and demonstrate that glucose re-directs IRF6 to genomic regions containing dimeric IRF-binding DNA motifs associated with active differentiation gene transcription.

To further characterize sites of glucose-modulated IRF6 action in the genome, chromatin accessibility in CRISPR-mediated IRF6 knockout differentiating keratinocytes was assessed using ATAC-seq (**Table S9**). IRF6-dependent accessible regions were overlapped with IRF6-regulated genes and IRF6 ChIP-seq data to nominate a subset of genes directly bound and regulated by IRF6 whose chromatin accessibility is also impacted by IRF6 (**Fig. 4g**). Among these putative glucose-dependent, IRF6 direct targets were key pro-differentiation TFs, including *GRHL1, GRHL3, HOPX* and *PRDM1*, which each play essential roles in epidermal differentiation. Expansion of IRF6 and glucose-regulated genes to those whose expression is downregulated with IRF6 depletion but retain accessible chromatin (stable ATAC-seq peaks), identified additional essential differentiation-related factors, notably, *PPARD, SBSN, CALML5* and *TGM1* (**Fig. 4h**). IRF6-bound distal H3K27ac-marked PIRs, representing putative enhancer elements looped to *HOPX, GRHL3* and *PRDM1* promoters in the presence of glucose; IRF6 enhancer binding was diminished, chromatin accessibility was decreased, and expression of these genes was reduced with glucose restriction (**Fig. 4i, Extended Data Fig. 8h-k**). IRF6 occupancy was not an obligate requirement for enhanced chromatin accessibility at all sites, however, as shown at the IRF6-dependent *SBSN* target gene locus (**Extended Data Fig. 8k**). Glucose therefore targets IRF6 to specific genomic sites where it may modulate chromatin accessibility and target gene expression. Taken together, these data support a working model (**Extended Data Fig. 9**) in which a tissue gradient of increasing glucose promotes differentiation by facilitating the dimerization and genomic binding of the IRF6 pro-differentiation TF. The mechanistic basis for such a glucose gradient within epidermis and whether similar gradients exist in other differentiating tissues represent intriguing topics for future study as does the potential for additional catabolism-independent roles for glucose as a modulator of protein multimerization. The 497 glucose-associated proteins identified here provide a resource for future efforts designed to explore the existence of additional glucose roles as a biomolecular modulator of diverse regulatory proteins.

## ACKNOWLEDGEMENTS

We thank J.Z. Long, H.Y. Chang, M.P. Snyder, W.J. Greenleaf, & A.E. Oro for pre-submission review. We thank B. Zarnegar, J. Kovalski, I. Alam, L. Looger, & J. Keller for helpful discussions & reagents, P. Neela for technical assistance, and A. Dazey and P. Bernstein for administrative assistance. This work was supported by the U.S. Department of Veterans Affairs Office of Research and Development (PAK), NIAMS R01 AR043799 and AR045192 (PAK) and K01 AR070895 (VLP) and in part by NIH P30 CA124435 utilizing the Stanford Cancer Institute Proteomics Shared Resource. This work is dedicated to the memory of coauthor S.S. Gambhir, whose untimely passing occurred during the completion of this work.

## AUTHOR CONTRIBUTIONS

Conceptualization: VLP, AB, and PAK, Formal analysis: VLP, AB, GG, YZ, LD, MGG, ZS, WM, ALJ, Methodology: VLP, AB and GG, Investigation: VLP, AB, AG, GG, LD, ZS, WM, DTN, AML, RLS, RMM, OSG, Resources: VLP, AB, GG, SSG, JY, PAK, Writing: VLP, AB and PAK, Funding Acquisition: PAK, Supervision: PAK.

## CONTACT FOR REAGENT AND RESOURCE SHARING

Further information and requests for resources and reagents should be directed to and will be fulfilled by the Lead Contact, Paul A. Khavari.

