## Supplemental Information for "Glucose modulates transcription factor dimerization to enable tissue differentiation"

**EXTENDED DATA FIGURE LEGENDS**

**Extended Data Figure 1. Metabolomic analysis identifies glucose accumulation in differentiated keratinocytes.** Enriched metabolic pathways in metabolomic data from 5 mass spectrometry methods, including UPLC-MS, positive and negative ionization LC-MS, LC-polar-MS, and GC-MS, in **(a)** day 3 (d3) differentiated keratinocytes or **(b)** day 6 (d6) differentiating keratinocytes. Levels of trans-urocanate **(c)** and glucose **(d)** from MS data in differentiating keratinocytes (n=5 biological replicates). **(e)** Quantitation of 2-NBDG fluorescence in progenitor (PG), d3 or d6 differentiated keratinocytes (n=3 biological replicates, 2 fields per replicate from Fig. 1b). **(f)** Quantitation of ^3^H-FDG accumulation in progenitor, d3 or d6 keratinocytes (n=3 biological replicates, 3 technical replicates each). **(g)** Western blot analysis of PG, d3 or d6 keratinocytes transduced with empty vector control or the glucose sensor, iGlucoSnFR-Myc. **(h)** Representative fluorescence images of keratinocytes transduced with empty vector or glucose sensor, iGlucoSnFR-Myc in PG or d7 differentiated keratinocytes grown in standard (4.5g/L) or low (0.5g/L) glucose media. GFP signal indicates detection of free glucose (n=2 biological replicates, scale bar=20μm). **(i)** Representative fluorescence image of keratinocytes transduced with empty vector or NLS-iGlucoSnFR-Myc in PG or d7 differentiated keratinocytes. GFP signal indicates detection of free glucose in the nucleus (n=2 biological replicates, scale bar=20μm). **(j)** Immunofluorescence staining of GFP [green] and nuclear DAPI stain [blue] in epidermal organoids (d7) generated from keratinocytes transduced with empty vector control or iGlucoSnFR-Myc; hematoxylin and eosin staining of epidermal organoids is also shown (n=2 biological replicates, A and B, scale bar=20μm). **(k)** Depiction of regions delimited for the stratum corneum/granular, spinous, or basal layers for GFP quantitation of epidermal organoids in Figure 1c. Because of their adjacency and thinness, the granular layer and stratum corneum are quantified together (n=3 biological replicates, two replicates are shown, scale bar=20μm). **(l)** 2-NBDG, a fluorescent glucose analog, accumulates in adipocytes, myoblasts, and osteoblasts; 2-NBDG [green], Hoechst nuclear stain [blue] (scale bar=10μm). **(m)** Quantitation of fluorescence in (l) (n=3 biological replicates and at least 4 fields per replicate). Quantitation of radiolabeled glucose ^14^C-Glucose (C6) accumulation after **(n)** 24h or **(o)** 96h, normalized to mg of protein in each sample (n=3 biological replicates and 3 technical replicates per experiment). **(p)** qPCR of differentiation markers specific for adipocytes, myoblasts or osteoblasts confirming differentiation of each cell type (n=2 biological replicates). Statistical significance determined using unpaired two-tailed Student *t*-test. Error bars represent S.E.M., *p<0.05, **p<0.01, ***p<0.005, ****p<0.001.

**Extended Data Figure 2. Glucose metabolism intermediates and outputs in epidermal differentiation.** Levels of metabolites related to the 3 major glucose catabolic pathways **(a)** glycolysis, **(b)** pentose phosphate, and **(c)** hexosamine from MS data in d3 or d6 differentiated keratinocytes (n=5 biological replicates). **(d)** Western blot analysis of the post-translational modification O-GlcNAc in progenitor (PG), d3 or d6 keratinocytes grown in standard (4.5g/L) or low (0.5g/L) glucose media. **(e)** Quantitation of immunoblot in (d). Metabolic flux analysis was conducted on progenitor (PG), d3 and d6 differentiated keratinocytes to evaluate [U-^13^C] glucose incorporation into **(f)** glucose-6-phosphate/glucose-1-phosphate/fructose-6-phosphate, **(g)** fructose-1,6-bisphosphate, **(h)** lactate, **(i)** pyruvate or **(j)** a-ketoglutarate (n=3 biological replicates per time point). **(k)** Levels of metabolites related to fatty acid metabolism from MS data in d3 or d6 differentiated keratinocytes (n=5 biological replicates). **(l)** Western blot analysis of a panel of lipid metabolism regulators in PG, d3 and d6 differentiated keratinocytes (n=3 biological replicates). Statistical significance determined using unpaired two-tailed Student *t*-test. Error bars represent S.E.M., *p<0.05, **p<0.01, ***p<0.005, ****p<0.001.

**Extended Data Figure 3. Impacts of glucose concentration on epidermal differentiation. (a)** Measurement of proliferation of progenitor keratinocytes in standard (4.5g/L), low (0.5g/L) or no glucose media (n=3 biological replicates). **(b)** Intracellular glucose levels of d3 differentiated keratinocytes grown in media with varying glucose concentrations (0, 0.5, 1.5, 2.5, 3.5 or 4.5g/L) (n=3 biological replicates). **(c)** Heatmap of qPCR analysis of epidermal differentiation gene expression in varying glucose concentration media shown in (b). **(d)** Immunostaining of differentiation markers [red], collagen VII [green], and nuclear DAPI stain [blue] in epidermal organoids generated with keratinocytes transduced with empty vector (EV) control or HA-hexokinase 1 and 2 (HK1/2) (n=2 biological replicates, scale bar=20μm). **(e)** Western blot analysis of keratinocytes transduced with EV control or HK1/2, along with quantitation of immunoblot in **(f)**. **(g)** Intracellular glucose levels in differentiated keratinocytes overexpressing HK1/2 (n=2 biological replicates). **(h)** qPCR of epidermal differentiation gene expression in epidermal organoids overexpressing HK1/2 (n=2 biological replicates). **(i)** qPCR of epidermal differentiation gene expression in differentiated keratinocytes (d3) transduced with EV control or HK1/2 treated with DMSO or 2-deoxy-d-glucose (2DG) (n=2 biological replicates). **(j)** Measurement of intracellular glucose levels in (i) (n=2 biological replicates). **(k)** Intracellular glucose levels in differentiated keratinocytes transduced with empty vector control or G6PD (n=2 biological replicates). **(l)** qPCR of epidermal differentiation gene expression in d3 differentiated keratinocytes transduced with EV control or G6PD (n=2 biological replicates). **(m)** qPCR of epidermal differentiation gene expression in d3 differentiated keratinocytes transduced with EV control or G6PD treated with DMSO or the DHEA inhibitor of G6PD (n=2 biological replicates). **(n)** Measurement of intracellular glucose levels in (m) (n=2 biological replicates). Statistical significance determined using unpaired two-tailed Student *t*-test. Error bars represent S.D., *p<0.05, **p<0.01, ***p<0.005, ****p<0.001.

**Extended Data Figure 4. Glucose transporters in epidermal differentiation.**

Western blot analysis of **(a)** GLUT1, **(b)** GLUT3 or **(c)** SGLT1 in progenitor (PG) and d3 differentiated keratinocytes (GLUT1 protein runs as a broad band on western analysis). Expression of **(d)** GLUT1 (*SLC2A1*) and **(e)** GLUT3 (*SLC2A3*) in single-cell RNA-seq data from normal human epidermis in the basal, cycling and differentiated cell populations. **(f)** Analysis of *SLC2A1*, *SLC2A3*, and *SLC5A1* regulation by epidermal transcription factors (TFs); gray represents gene expression downregulation with TF knockdown or knockout in keratinocytes interrogated in publicly available data compiled in Lopez-Pajares et al. **(g)** Immunofluorescence staining of glucose transporters [red] and nuclear DAPI stain [blue] in PG and d3 differentiated keratinocytes (scale bar=10μm). **(h)** Western blot analysis of GLUT1 protein levels in keratinocytes treated with non-targeting (NT) or sgRNAs targeting *SLC2A1* (GLUT1). **(i)** qPCR of differentiation gene expression in *SLC2A1* KO d3 differentiated keratinocytes (n=2 biological replicates). **(j)** Western blot analysis of GLUT3 protein levels in keratinocytes treated with NT or sgRNAs targeting *SLC2A3* (GLUT3). **(k)** qPCR of differentiation gene expression in *SLC2A3* KO d3 differentiated keratinocytes (n=2 biological replicates). **(l)** qPCR of differentiation gene expression in *SLC5A1* (SGLT1) KO keratinocytes (n=2 biological replicates). **(m)** Relative intracellular glucose concentrations in differentiated keratinocytes treated with DMSO, GLUT inhibitor WZB117 (10μM), or SGLT inhibitor phlorizin (50μM) (n=3 biological replicates). **(n)** qPCR analysis of differentiation gene expression in differentiated keratinocytes treated with DMSO, WZB117 or phlorizin (n=3 biological replicates). Statistical significance determined using unpaired two-tailed Student *t*-test. Error bars represent S.E.M., *p<0.05, **p<0.01, ***p<0.005.

**Extended Data Figure 5. Identification of IRF6 as a glucose-binding pro-differentiation transcription factor. (a)** Top 10 terms for pathway enrichment using the BioPlanet database for 497 glucose-binding proteins (Enrichr analysis). **(b)** Heatmap of 497 proteins identified as glucose-binding proteins showing mRNA expression from RNA-seq in early (d3) and late (d6) differentiated keratinocytes, and enrichment in glucose elution versus galactose elution in dextran column affinity chromatography followed by LC-MS/MS. **(c)** Enriched GO terms for glucose-regulated genes with open (increased) ATAC-seq peaks. **(d)** Epidermal organoids of primary human keratinocytes transduced with Cas9 and non-targeting (NT) control sgRNA or *IRF6*-targeting sgRNA representing IRF6 knock-out (KO) human tissue harvested at d7; differentiation markers [red], nuclei [blue], collagen VII basement membrane staining [green] (n=3 biological replicates, representative image shown, scale bar=20μm). **(e)** Western blot analysis of keratinocytes transduced with Cas9 and control NT sgRNA or 4 different *IRF6* targeting sgRNAs, used to generate epidermal organoids. **(f)** Quantitation of IRF6 protein levels in (e). **(g)** Differentiation gene expression in *IRF6* KO tissue measured by qPCR (n=2 biological replicates per sample). **(h)** Volcano plot of PAS-seq data from *IRF6* KO tissue showing 720 upregulated and 1191 downregulated genes (n=3 biological replicates, FDR>0.05, fold change>1.5). Highlighted are downregulated differentiation-specific genes. **(i)** Enriched biological process GO terms for *IRF6* KO tissue downregulated genes. **(j)** Overlap of *IRF6* KO downregulated genes with glucose-regulated genes and the epidermal differentiation gene signatures generated by Lopez-Pajares et al. and Rubin et al. Statistical significance determined using unpaired two-tailed Student *t*-test. Error bars represent S.D., *p<0.05, **p<0.01, ***p<0.005, ****p<0.001.

**Extended Data Figure 6. IRF6 glucose binding capacity is required for epidermal differentiation.** **(a)** Microscale thermophoresis (MST) analysis of IRF6 affinity toward the metabolically stable glucose analog, 3-O-methyl-glucose (K_d_=115±78.7μM) (n=3). **(b)** Quantitation of intracellular glucose levels in progenitor and d3 differentiated keratinocytes; (n=5 biological replicates). **(c)** Candidate IRF6 peptides enriched for glucose binding (n=2). **(d)** MS/MS of the IRF6 peptide identifying the P104 and N106 residue crosslinked to glucose. **(e)** Modeling the IRF6-glucose interaction using Autodock Vina, showing a highly scored model (score<-5) and the location of the unique peptide identified via azido-glucose IP-MS (blue) and peptides identified via UV-C crosslinking of glucose to IRF6 followed by MS (orange); the enlarged image shows the location of disease-associated residue arginine 84 (purple), as well as proline 104 (green) and asparagine 106 (cyan) identified as glucose-binding residues. **(f)** Electrophoretic mobility shift assay of recombinant IRF6 and P104G or N106A glucose-binding deficient mutants. **(g)** Gene expression of d4 differentiated keratinocytes treated with control or *IRF6* siRNA with enforced expression of Flag-HA-6XHis (FHH) tagged-WT, P104G, N106A or P104G/N106A IRF6 constructs to rescue the *IRF6i* differentiation defect (representative technical replicates shown, n=2 biological replicates). **(h)** Western blot analysis of samples in (g). **(i)** Sanger sequencing chromatogram of targeted editing sequence at the endogenous *IRF6* gene locus; editing efficiencies are noted for each of the two replicates (n=2 biological replicates). **(j)** Immunostaining of keratin 10 [green], transglutaminase [red], and nuclear DAPI [blue] in epidermal organoids generated from R84R and R84C gene edited keratinocytes; (shown is a representative image from 2 biological replicates, scale bar=20μm). **(k)** Gene expression analysis of R84R and R84C gene edited tissue (n=2 biological replicates). **(l)** Immunostaining of keratin 10 [green], transglutaminase [red], and nuclear DAPI [blue] in epidermal organoids generated from P104P and P104G gene edited keratinocytes (shown is a representative image from 2 biological replicates, scale bar=20μm). **(m)** Gene expression analysis of P104P and P104G gene edited tissue (n=2 biological replicates). Statistical significance determined using unpaired two-tailed Student *t*-test. Error bars represent S.E.M., *p<0.05, **p<0.01, ***p<0.005, ****p<0.001.

**Extended Data Figure 7. Functional impacts of glucose on IRF6. (a)** Affinity of IRF6 towards cognate DNA binding sequence in the *SFN* locus measured by MST. Measurements were conducted using a wild-type *SFN* sequence DNA oligo in PBS (K_d_=16.4±5.8μM) or 350μM glucose (K_d_=0.357±0.215μM) (n=4). **(b)** Summary of the change in affinity of IRF6 toward DNA ± 350μM glucose, log scale shown. **(c)** Affinity of MAFB towards cognate DNA binding sequence in the *GRHL3* locus measured by MST. Measurements were conducted using a wild-type *GRHL3* sequence oligo in PBS (K_d_=14.3±0.66μM) or 350μM glucose (K_d_=17.5±1.19μM) (n=3). **(d)** Electrophoretic mobility shift assay of IRF6 motif probe incubated with recombinant IRF6 with or without glucose or incubation with scrambled or competing oligonucleotides. Indicated are the free DNA and IRF6-DNA complex. **(e)** DNA ELISA of recombinant IRF6 binding to an IRF responsive element with or without glucose (n=2). **(f)** Western blot analysis of replicate samples of FHH-IRF6 co-immunoprecipitation from FHH-IRF6 and V5-IRF6 transduced keratinocytes differentiated for 3 days in standard (4.5g/L) or low (0.5g/L) glucose media and quantitation **(g)** (n=2 biological replicates). **(h)** Western blot analysis of FHH-IRF6 co-immunoprecipitation from FHH-IRF6 and V5-IRF6 transduced progenitor keratinocytes in cytoplasmic or nuclear cellular fractions. **(i)** Western blot analysis of FHH-IRF6 or FHH-IRF6^S413A/S424A^ mutant (S/A) and V5-IRF6 transduced keratinocytes differentiated in standard or low glucose media for 3 days after FLAG immunoprecipitation. **(j)** IRF6 luciferase reporter assay for d3 differentiated keratinocytes grown in standard or low glucose media (n=4 biological replicates). **(k)** Western blot analysis of cellular fractionation of d3 differentiated keratinocytes grown in standard or low glucose media along with **(l)** quantitation of IRF6 protein levels in nuclear extracts. Statistical significance determined using unpaired two-tailed Student *t*-test. Error bars represent S.E.M., *p<0.05.

**Extended Data Figure 8. Glucose-mediated IRF6 gene regulation. (a)** Normalized enrichment score for IRF6 ChIP-seq peaks in differentiated keratinocytes grown in standard or low glucose media (n=2 biological replicates). **(b)** Overlap of genes connected (cGenes) to IRF6 ChIP-seq peaks in standard (3858 genes) or low (1988 genes) glucose media (n=2 biological replicates). **(c)** HOMER de novo motif analysis of shared IRF6 ChIP-seq peaks in standard and low glucose media compared to known IRF6 JASPAR motif 2. **(d)** Gene ontology (GO) terms for glucose-modulated IRF6 ChIP-seq peaks identified in standard or low glucose conditions. **(e)** Top three enriched HOMER de novo motifs for 916 glucose-regulated ChIP-seq peaks. **(f)** Genomic features associated with standard, low or glucose-regulated IRF6 ChIP-seq peaks. **(g)** Location of IRF6 binding relative to TSS in standard, low or glucose-regulated ChIP-seq peaks. **(h)** IRF6 chromatin immunoprecipitation followed by qPCR of the *DMKN*, *SFN,* *GRHL3* and *HOPX* loci after d3 differentiation in standard glucose media (4.5g/L), standard glucose media with GLUT and SGLT1 inhibitors (10μM WZB-117 and 50μM phlorizin) treated for 15 h, or low glucose media (0.5g/L) (representative data shown is 2 technical replicates, n=2 biological replicates). Genome browser tracks of IRF6 ChIP-seq peaks and ATAC-seq peaks in d3 differentiated keratinocytes in standard or low glucose media, and ATAC-seq peaks in control (NT) or *IRF6* KO keratinocytes for the **(i)** *GRHL3*, **(j)** *PRDM1*, and **(k)** *SBSN/DMKN* loci. Error bars represent S.E.M.

**Extended Data Figure 9. Model of glucose-mediated gene regulation in epidermal differentiation.** Glucose accumulation is a feature of epidermal differentiation and free glucose accumulates in the differentiated layers of the epidermis. In progenitor keratinocytes, glucose equilibrium is maintained by the facilitative glucose transporters, GLUT1 and GLUT3 and glucose is metabolized. Upon differentiation, the sodium-coupled glucose co-transporter, SGLT1, is induced and glucose accumulation occurs, without increased glucose catabolism. Glucose binds the IRF6 pro-differentiation TF. Glucose enhances IRF6 dimerization and genomic targeting to regulate gene expression of IRF6-dependent target genes, including pro-differentiation transcription factors.

**SUPPLEMENTARY TABLE TITLES**

**Table S1. Metabolomic analysis of progenitor and differentiated keratinocytes.**

**Table S2. PAS-seq analysis of differentially expressed genes in standard vs low glucose epidermal organoids.**

**Table S3. List of putative transcription factors regulating *SLC2A1*, *SLC2A3* and *SLC5A1* predicted using ATAC-seq foot printing analysis.**

**Table S4. List of proteins identified in dextran glucose polymer column affinity purification mass spectrometry with enrichment over galactose.**

**Table S5. Connected genes (cGenes) assigned to increased/open ATAC-seq peaks in standard or low glucose or IRF6 KO differentiated keratinocytes.**

**Table S6. PAS-seq analysis of differentially expressed genes in IRF6 KO epidermal organoids.**

**Table S7. IRF6 azido-glucose MS and crosslinking MS.**

**Table S8. IRF6 ChIP-seq peaks and cGenes in standard versus low glucose conditions.**

**Table S9. Connected genes (cGenes) assigned to decreased/closed IRF6 KO ATAC-seq peaks in differentiated keratinocytes.**

**Table S10. Sequences for sgRNAs used for CRISPR-Cas9 KO.**

**Table S11. Sequences for primers used in RT-qPCR analysis.**

**Table S12. Sequences for DNA oligos used in MST analysis.**

**METHODS**

**Human keratinocyte isolation and culture**

Primary human keratinocytes were isolated from fresh, surgically discarded neonatal foreskin. All human cells were collected and analyzed by protocols approved by the Stanford Human Subjects Institutional Review Board and in accordance with the NIH genomic data sharing policy. Keratinocytes were maintained in a 1:1 mixture of Keratinocyte-SFM (Life Technologies #17005-142) and Medium 154 (Life Technologies #M-154-500). Keratinocytes were maintained by growth at sub-confluence or seeded for differentiation at full confluence and grown for 3 or 6 days or seeded onto dermis to generate epidermal organoids. Differentiation media consisted of DMEM, no glucose (Thermo Fisher) supplemented with sodium bicarbonate (Thermo Fisher), keratinocyte-SFM supplements (Thermo Fisher), glucose (Sigma) and 1.2mM calcium (Sigma).

**Metabolomics**

Metabolomics analysis was performed in sub-confluent, day 3 and day 6 differentiated keratinocytes. These experiments were performed on 5 biological replicates, and were conducted by Metabolon, Inc. Five mass spectrometry methods, including UPLC-MS, positive and negative ionization LC-MS, LC-polar-MS, and GC-MS were used to identify 614 compounds in these keratinocyte cells.

**Glucose detection and tracing**

2-NBDG was incubated with cells for 3 hours at a concentration of 250μM. Prior to imaging, cells were washed following manufacturer instructions (Life Technologies N13195). Images were taken and processed using a Zeiss Axio Observer Z1 microscope with ApoTome.2 and Zeiss Axiovision software. The iGlucoSnFR glucose sensor ^1^ was cloned into a lentiviral expression vector (pLEX) and transduced into human keratinocytes. After selection with puromycin, cells were seeded at sub-confluence or full confluence to initiate differentiation, or seeded on human dermis to generate epidermal organoids. Cells were tracked by live cell imaging in 2D culture using GFP. For organoids, samples were harvest at d7 and fixed in 4% formalin. Cryosections were directly visualized for GFP fluorescence or stained for differentiation markers and GFP and was visualized under fluorescence microscopy. Hematoxylin and eosin staining was performed according to the manufacturer instructions (Abcam). Radiolabeled glucose tracers were incubated with cells for the amount of time specified in each experiment at a concentration of 0.1 microCurie/500 microliters. After incubation, cells were washed twice with cold PBS and lysed with modified RIPA buffer [50 mM Tris pH 7.3, 0.5% Sodium Deoxycholate, 1% NP-40, 0.01% SDS, 150 mM NaCl] supplemented with EDTA-free protease inhibitor cocktail tablets (Pierce). 100 μL was used for scintillation counting and 10 μL was used in duplicate for total protein BCA measurements (Thermo Fisher Scientific 23225). A standard of the media with tracer was also quantitated by scintillation counting and used to calculate the total activity present in each well to enable percent uptake and percent accumulation calculations. Pulse chase and export experiments were performed similarly, except after 5 minutes of incubation, cells were washed twice with room temperature PBS and incubated for time indicated in standard media without tracer. Cells and media were analyzed by scintillation counting.

**Metabolic Measurements**

The extracellular acidification rate (ECAR) and oxygen consumption were measured with the Seahorse XF24 analyzer. In all cases 3 biological replicates were performed.

**Metabolic tracing**

For progenitor keratinocytes, cells were plated at a density of 500,000 cells in 60 mm dishes, and differentiated keratinocytes were plated at a density of 2 million cells per 60 mm dishes in glucose-free DMEM (Thermo Fisher) supplemented with sodium pyruvate (Thermo Fisher), keratinocyte-SFM supplements (Thermo Fisher), 4.5g/L glucose (Sigma) and antibiotics (Thermo Fisher). The following was performed 24h after seeding progenitors or at d3 or d6 of differentiation: media was replaced with glucose-free DMEM (Thermo Fisher) and 4.5g/L ^13^C_6_-glucose (Sigma). Media was switched to glucose-containing media 30 min, 1 hr, 2 hr, 6hr, and 24 hr prior to analysis (n = 3 per group).

**Metabolite harvesting and liquid chromatography-mass spectrometry analysis**

Cells were washed with cold PBS, lysed in 80% Ultra LC-MS acetonitrile (Thermo Scientific) on ice for 15 minutes, and centrifuged for 10 minutes at 20,000 x g at 4 °C. 200 μL of supernatants were subjected to mass spectrometry analysis. Liquid chromatography was performed using an Agilent 1290 Infinity LC system (Agilent, Santa Clara, US) coupled to a Q-TOF 6545 mass spectrometer (Agilent, Santa Clara, US). A hydrophilic interaction chromatography (HILIC) method with a BEH amide column (150 x 2.1 mm, 1.7 μm; Waters) was used for compound separation at 35 °C with a flow rate of 0.3 mL/min. Mobile phase A consisted of 20 mM ammonium acetate and 20 mM ammonium hydroxide in water and mobile phase B was acetonitrile. The gradient elution was 0—1 min, 85% B; 1—12 min, 85% B → 65% B; 12—12.2 min, 65% B → 40% B, 12.2-15 min, 40% B. After the gradient, the column was re-equilibrated at 85% B for 5 min. The overall runtime was 20 minutes, and the injection volume was 5 μL. The Agilent Q-TOF was operated in negative mode and the relevant parameters were as listed: ion spray voltage, 3500 V; nozzle voltage, 1000 V; fragmentor voltage, 125 V; drying gas flow, 11 L/min; capillary temperature, 325 °C; drying gas temperature, 350 °C; and nebulizer pressure, 40 psi. A full scan range was set at 50 to 1600 (m/z). The reference masses were 119.0363 and 980.0164. The acquisition rate was 2 spectra/s. Targeted analysis, isotopologues extraction (for the metabolic tracing study), and natural isotope abundance correction were performed by the Agilent Profinder B.10.00 Software (Agilent Technologies).

**Epidermal Organoids**

Primary human keratinocytes were isolated from fresh surgically discarded skin and cultured in a 1:1 mix of Keratinocyte-SFM (Life Technologies #17005-142) and Medium 154 (Life Technologies #M-154-500). Organotypic regeneration of human epidermis was performed as previously described ^2^. Briefly, 500,000 cells were seeded onto devitalized human dermis in keratinocyte growth media (KGM) at the air-liquid interface and allowed to grow for specified amount of time (3-7 days).

**Immunostaining and immunofluorescent microscopy**

For immunofluorescence staining, tissue sections (7 μm thick) were fixed using either acetone, methanol or 4% paraformaldehyde (Pierce). Primary antibodies were incubated at 4°C overnight, and secondary antibodies were incubated at room temperature for 1 hour. Slides were mounted using Duolink In Situ Mounting Medium with DAPI (Sigma, DUO82040). All images were taken and processed using a Zeiss Axio Observer Z1 microscope with ApoTome.2 and Zeiss Axiovision software. For quantitation, ImageJ software was used. The antibodies used in this study were Rb-anti-HA (CST, #3724), anti-KRT1 (Covance, PRB-149P), anti-KRT10 (Neomarkers, MS611P), Ms-anti-CollagenVII (Chemicon, MAB1345), pAb-anti-CollagenVII (Chemicon, 234192), anti-Filaggrin (Abcam, ab218863), anti-Loricrin (Covance, PRB-145P), anti-Transglutaminase (Biomedical Technology, BT-621).

**Gene transfer and CRISPR/Cas9 knockout**

Gene transfer was performed by viral transduction. Gene cDNA was cloned into pLEX lentiviral vector. IRF6 cDNA was acquired from Origene (SC116274). The R84C and S413/424A mutations were introduced using PCR mutagenesis using the InFusion kit (Takara). Lentivirus was produced in HEK293T Lenti-X cells (Takara) seeded in 10cm plates at 70% confluence. Cells in 10cm were transfected with 6 μg of p8.91, 2 μg of pMDG, and 8 μg of gene of interest in 1.2mL Opti-MEM (Thermo Fisher Scientific) and 45μL of Lipofectamine 2000 (Thermo Fisher Scientific). Media was replaced after 15 hours. Supernatant was collected 48 hours after transfection, filtered through a 0.45 um PES membrane, and concentrated to 50X with Lenti-X concentrator (Takara). Human keratinocytes were transduced with pLEX-Cas9 lentivirus (Addgene: 117987) ^3^ and single guide RNA virus and selected with blasticidin-HCl and puromycin for 48h. Cas9 expression was confirmed by anti-FLAG (F3165, Millipore Sigma) western blotting, sgRNA expression was verified by mCherry visualization and knockout was verified by qPCR and/or Western blot analysis. The sgRNA sequences are listed in table S9.

CRISPR-Cas9/AAV gene editing vector construction and virus production

For each editing experiment two consecutive ssAAV vectors were constructed differed by presence of the mutation of interest or a synonymous substitution control. The donor ssAAV vector used as a template for homologous directed recombination (HDR) contained ~1000bp genomic DNA sequence flanking the 5’ and 3’ end of the target mutations between AAV ITR sequences. First, 811nt flanking 5’ end of the *IRF6* intron/exon 4 junction was amplified using primers IRFF and IRFR and In-Fusion assembled into NheI/EcoRI digested AAV transfer plasmid following with In-Fusion assembly of the 1005nt gBlock gene fragments (IDT) containing *IRF6* exon 4 and 3’ end of the intron with mutations of interest cloned into StuI digested AAV recombinant vector. The following point mutations were engineered within the gBlock gene fragments: R84R (CGC/CGG), R84C (CGC/TGC), P104P (CCC/CCT), P104G (CCC/GGC), N106N (AAC/AAT) and N106A (AAC/GCC). Additionally, in order to prevent endonuclease mediated re-cutting, PAM mutations were introduced resulting to the synonymous amino acid substitution: G/A (K80K) and G/A (E202E). For genomic DNA amplification, following primers were used:

IRFF: 5’ ATCAACGCGTGCTAGCTTACTCTTGTTGCCCAGGCTC

IRFR: 5’ GCTTGATATCGAATTAGGCCTGAGAACAAGAAACCAC

After confirmation of the insert sequence integrity, constructs were used for AAV virus production using “Helper-free” AAV system consist of pHelper, pAAV-DJ plasmids and 293AAV cell line (Cell Biolabs). Briefly, 50–70% confluent 293AAV cells grown in DMEM supplemented with 10% FBS were triple transfected with pHelper, pAAV ITR-transfer, and pAAV Rep-Cap plasmids using FuGENE6 (Promega) reagent in two 10cm dishes following manufacturer’s recommendations. After transfection, medium changes to DMEM with FBS were performed in 24hrs. At day 3 post-transfection, media and cells were collected and processed separately. Cells were lysed in lyses buffer (20mM Tris pH8.0, 150mM NaCl, 2mM MgCl_2_) via freeze thaw and the homogenates were cleared from debris by centrifugation. AAVs were precipitated from medium with polyethylene glycol (PEG) 8000. The PEG-precipitated AAV was collected by centrifugation, resuspended in lyses buffer and combined with cell lysate fraction. The crude virus was tittered^4^ and AAV-DJ serotyped donor ssAAV virus was produced at genomic titer of 1.5-5x10^13^ TU/mL and used for HDR experiments at MOI 2.5x10^5^.

***CRISPR and AAV mediated HDR***

The guide sequence targeting *IRF6* for CRISPR/Cas9 genome editing was predicted using CHOPCHOP web tool^5^ and were ordered from IDT as sgRNA accordingly. Following gRNAs were used in this study: gRNA IRF6A: 5’ GATGTATGATGGCACCAAGG, gRNAIRF6B: 5’ CCCTGACCCAGCTAAATGGA. For CRISPR/Cas9 genome editing 73 pmol of the sgRNA was complexed with 61 pmol recombinant Alt-RspCas9 protein (IDT) in 10ml of Amaxa nucleofection buffer for keratinocytes (Lonza) for 10min and immediately used for nucleofection of 8x10^5^ primary keratinocytes with Amaxa nucleofection apparatus (Lonza) using program T-018. After recovery cells were mixed with AAV virus containing either wt or mutant donor template at MOI 2.5x10^5^, split into 2 wells of a 6-well plate and propagated for 72hrs. After culture reached 50-60% confluence cells were grown into 10 cm plates during which genomic DNA was isolated and evaluated for the editing efficiency using PCR amplification and sequencing of the bulk cell papulation, as well as by cloning of the amplified fragment into pBluescript vector and evaluating editing efficiency by individual colony sequencing. The population of cells with at least over 80% editing efficiency were considered for further experiments.

**Glucose transporter inhibition**

The WZB-117 (Sigma) inhibitor targeting GLUT1 and GLUT3 was applied to media of differentiated keratinocytes at a concentration of 10uM for 24h prior to harvesting. The SGLT1 inhibitor, phlorizin (Sigma) was used at a concentration of 50uM for 24h prior to harvesting.

**RNA extraction**

RNA was extracted using the RNeasy kit (Qiagen). For 2D culture, cells were lysed directly in RLT buffer and for epidermal organoids, tissue was harvested, and cells were scraped off of the dermis then placed in RLT buffer.

**RT-qPCR**

Reverse transcription was performed using 1μg of RNA extracted from cells or tissue using the iScript cDNA synthesis kit (Bio-rad). RT-qPCR was performed using the 2X Maxima SYBR Green/ROX qPCR Master Mix (ThermoFisher). Table S10 contains sequences for primers used in RT-qPCR analysis.

**Electrophoretic Mobility Shift Assay**

The probe was synthesized with an oligonucleotide sequence containing a trimer of the IRF6 DNA binding motif with a modification at the 5’ end containing IRD700 (IDT): 5’-TGGTTTCGGTTGACTTGGTTTCGGTTGACTTGGTTTCGGTTGA-3’. Oligos were annealed and diluted to a concentration of 50nM. “Cold” oligonucleotides did not contain the IRD700 modification and consisted of the sequence noted above or the following sequence for the non-competing/mutant oligo:

5’-ACGACCAACTCAATCACGACCAACTCAATCACGACCAACTCAA-3’ and were diluted to a concentration of 5uM. For the reactions, 1ul of probe was incubated at room temperature for 30 min with IRF6 recombinant protein in reaction buffer [20mM Tris, pH 7.5, 50mM NaCl, 1mM EDTA, 5% glycerol, 5mM dithiothreitol, 1ug bovine serum albumin, 100ng poly (dI:dI)-poly (dI:dC), 0.25% Tween-20] with or without 1ul of competing or mutant probe. Where indicated, 350uM glucose was added to the reaction. DNA-protein complexes were resolved on a 6% DNA retardation gel (Thermo Fisher) in 0.5X TBE for 1.5h at 100 V. The resulting gel was scanned on the Odyssey instrument (Li-COR) to detect IR700 and ImageStudio Lite software was used to visualize the DNA-protein complexes (Li-COR).

**ELISA**

The commercially available TransAM-IRF-3 DNA binding ELISA (Active Motif) was adapted for use with IRF6, as the kit is designed to assay DNA binding to the IRF core motif containing a full-site which IRF6 can bind to. The assay was performed according to the manufacturer’s instruction with following modifications: the IRF-3 antibody was replaced with an IRF6 goat antibody (Novus) and the secondary antibody was replaced with an HRP-conjugated anti-goat secondary antibody (Thermo Fisher).

**Luciferase Reporter Assay**

The IRF6 DNA binding sequence^6^: 5’-AATTCTCAACCGAAACTAGTCAACCGAAACTAGTCAACCGAAACTAGTCAACCGAAACTAGTCAACCGAAACTAGG-3’ or a scrambled control 5’-AATTCGAAACCGACTTATATCACAACGACCAAAATAAGCATCGCTCAATGGAACCCTCCCATACAGAGGAAACAAG-3’ was cloned into the pGreenfire lentiviral vector (System Biosciences). Lentivirus was generated in 293T cells and keratinocytes were transduced as noted above. Keratinocytes were seeded at full confluence; media was supplemented with 1.2mM calcium to induce differentiation and cells were harvested at d3. Cells were lysed directly in passive lysis buffer and the Luciferase Assay Reagent kit (Promega) was used per manufacturers protocol to detect firefly luciferase activity. A portion of cell lysate was retained for protein quantitation and luciferase signal was normalized to total protein amounts.

**Glucose affinity mass spectrometry**

***Dextran column***

Keratinocytes were differentiated by seeding at full confluence and harvested 3 days after seeding. Three 10cm plates of differentiated keratinocytes were lysed in RIPA buffer [0.1% SDS, 1% NP-40, 1% sodium deoxycholate, 1mM EDTA, protease inhibitors]. Dextran columns containing 5ml of Sephadex G100 resin were washed with 10ml of PBS and cell lysate was applied to the columns. Columns were washed 3 time with wash buffer [500mM Tris pH 7, 1M NaCl]. Columns were then eluted with 1ml of 1M glucose or 1M galactose. Samples were run on SDS-PAGE gel and a single gel slice was cut out. Samples were reduced in-gel with dithiothreitol, and alkylated with iodoacetamide. Processed proteins were subsequently digested in-gel, overnight, with Trypsin/Lys-C (Promega, V5071) at an enzyme/substrate ratio of 1:100 in 50 mM NH_4_HCO_3_ (pH 8.5) at 37°C. The next day, peptides were recovered from gels first using a solution containing 5% acetic acid in H_2_O and second with a solution containing 2.5% acetic acid in an equal volume mixture of CH_3_CN and H_2_O. The peptide mixture was subsequently dried in a Speed-vac and desalted using OMIX C18 pipet tips (Agilent Technologies, Santa Clara, CA). The timsTOFpro was operated in PASEF (parallel accumulation sequential fragmentation) mode, with mass tolerances of 50 ppm for both precursor and fragment ions. The Orbitrap fusion was operated with either collision-induced dissociation (CID) or electron transfer dissociation / higher-energy collisional dislocation (ETD/HCD) in a decision tree format for fragmentation to identify precursor peptides. Precursor and HCD were detected in the Orbitrap with 12 ppm mass tolerances, while CID and ETD were detected in the ion trap with 0.4 Da mass tolerances.

***Azido-glucose pulldown***

One million keratinocytes were seeded for differentiation at full confluence in 6-well plates and grown for 2.5 days at which point media was changed from standard glucose (4.5g/L) DMEM to a mix of 2.25g/L 2-azido-2-deoxy-D-glucose and 2.25g/L D-glucose DMEM and incubated for 15 hours. Cells were washed with PBS twice, crosslinked with 3000 joules of UV-C power then lysed in urea lysis buffer [200mM Tris pH8, 4% CHAPS, 1M NaCl, 8M Urea]. Following lysis, the Click-&-Go Click Chemistry Capture Kit (Click Chemistry Tools) was used to isolate azido-glucose linked proteins according to the manufacturer’s instructions. Samples were digested on-bead per the kit protocol and sent for LC-MS/MS analysis.

**Crosslinking mass spectrometry**

Human recombinant IRF6 (Origene) was diluted in PBS with or without 350μM glucose (Sigma) and incubated for 15 min at room temperature. The BS3 crosslinker was added at 0.2 or 0.5mM (Thermo Fisher Scientific) as previously described ^7^. The mixture was incubated for 1 h at 25°C at 350 rpm in a thermomixer. The reaction was quenched with the addition of 4X LDS, 10% beta-mercaptoethanol and heating at 75°C for 15 min. Samples were run on SDS-PAGE gel and the protein gels were stained with colloidal Instant Blue stain (Pierce) and the region containing the crosslinked complex was cut out, processed as described above, and sent for LC-MS/MS analysis.

**UV-C crosslinking mass spectrometry**

IRF6 recombinant protein (30 or 90 ng/µL) was incubated with or without 350 µM glucose at room temperature for 15 minutes and was crosslinked by UVC (254 nm, UV Stratalinker 2400) at 0.3 J/cm^2^ in 3 µL droplets on a parafilm, cross-linking a total of 15 droplets per sample. All 15 droplets for each sample were combined for in-gel digestion. Samples were loaded onto a 10% SDS-PAGE gel. After electrophoresis, the ~60kD IRF6 bands were cut into slices, reduced in-gel with dithiothreitol, and alkylated with iodoacetamide. Processed proteins were subsequently digested in-gel, overnight, with Trypsin/Lys-C (Promega, V5071) at an enzyme/substrate ratio of 1:100 in 50 mM NH_4_HCO_3_ (pH 8.5) at 37°C. The next day, peptides were recovered from gels first using a solution containing 5% acetic acid in H_2_O and second with a solution containing 2.5% acetic acid in an equal volume mixture of CH_3_CN and H_2_O. All the resulting peptide mixture was subsequently dried in a Speed-vac and desalted using OMIX C18 pipet tips (Agilent Technologies, A57003100). LC-MS/MS experiments were conducted on a Orbitrap Fusion nanoLC/MS instrument (Thermo Fisher).

**Recombinant protein purification**

293T cells were seeded at 70% confluence in 10cm plates and transfected with plasmids encoding desired FLAG-HA-6XHIS-tagged recombinant proteins in pLEX vector using Lipofectamine 2000 (ThermoFisher). Media was changed 15-18h post-transfection and cells were harvested 48h post-transfection. Cells were lysed in FLAG IP lysis buffer [50mM Tris, pH7.4, 150mM NaCl, 1mM EDTA, 1% Triton X-100, protease inhibitors] for 30 min. Cell lysates were applied to M2 (FLAG) resin in 15ml tubes per the manufacturer’s protocol for Batch FLAG protein purification (Sigma) and incubated for 4 hours. Resin/lysate mixture was applied to chromatography columns (Bio-Rad), washed with 20 bed-volumes of wash buffer [50mM Tris, pH 7.4, 500mM NaCl, 3mM EDTA, 0.5% NP-40, 0.1mM DTT, protease inhibitors] and eluted in 5ml of elution buffer [PBS, 0.02% Tween, 0.5mg/ml FLAG peptide]. Eluted proteins were applied to a protein concentration column (Millipore) and concentrated 50X. An aliquot of purified recombinant protein was run on an SDS-PAGE gel with BSA standards to determine protein concentrations. Gel was stained with InstantBlue colloidal blue stain (Sigma) for 1 hour, de-stained overnight in water and imaged on the Li-COR Odyssey machine (Li-COR Biosciences). ImageStudio Lite software (Li-COR Biosciences) was used to quantitate BSA protein standards and recombinant protein to determine the protein concentration. Protein quality was also assessed using the Tycho NT.6 instrument (Nanotemper) which uses label-free measurements of structural integrity (protein folding) by applying a temperature gradient to the sample.

**Microscale thermophoresis**

His-tagged proteins were labeled with His-tag labeling kit RED-tris-NTA 2^nd^ generation (Nanotemper Technologies) according to manufacturer’s instructions at a 1:2 dye to protein ratio. Briefly, 200nM recombinant protein was diluted in PBS-T and incubated with 5nM Red-tris-NTA dye for 1 hour. The level of label signal was checked via a capillary scan at 60% LED power and diluted to 300 units using the provided 1x PBS-T, resulting in a target protein concentration of 25nM. Protein was then incubated with either PBS or glucose for glucose-affinity measurements with indicated concentrations (total of 16 2-fold dilutions per run). For protein:DNA affinity measurements, 50nM protein was incubated with varying amounts of 80-100bp DNA oligonucleotides for 20 min at room temperature in the presence of poly I/C, to reduce background, and PBS or 350μM glucose. For protein:protein affinity measurements, human recombinant IRF6 (Origene) with no tag was incubated with His-tagged and labeled IRF6 in the presence of PBS or 350μM glucose for 10 min. Microscale thermophoresis measurements were taken on the Monolith NT.115 instrument using premium capillaries (Nanotemper Technologies). Analysis was conducted using Nanotemper software. Sequences for DNA oligos were used for protein:DNA MST assays are listed in Table S11.

**In silico small molecule interaction modeling**

Autodock Vina ^8^ docking module using Chimera software ^9^ was utilized to model the interaction between glucose and IRF6. The input structure for IRF6 was acquired from the Alpha-Fold modeled structure for IRF6 (AF-O14896-F1). The entire IRF6 molecule was selected to identify potential docking sites for glucose with the top scoring hits having a score value of -4.8.

**Western blot**

Cells were lysed in RIPA buffer [0.1% SDS, 1% NP-40, 1% Sodium deoxycholate, 1mM EDTA in PBS]. Lysates were incubated on ice for 30 min and supernatant was cleared by centrifugation at max speed. Protein quantitation was performed using the BCA assay kit (Promega). 10-20 μg of protein were loaded on an SDS-PAGE gel and transferred onto nitrocellulose membrane (Bio-Rad). Blots were probed with the following antibodies: anti-IRF6 (Cell Signaling), anti-FLAG (Sigma), anti-V5 (Cell Signaling), anti-HA (Abcam), anti-Myc (Abcam, ab32), anti-beta-actin (Sigma), anti-lamin A/C (Cell Signaling), anti-GLUT1 (Abcam), anti-GLUT3 (Abcam), anti-SGLT1 (Abcam), anti-HK1 (Abcam), anti-tubulin (Abcam), anti-O-GlcNAc (Cell Signaling). IR-dye conjugated secondary antibodies were used for detection on the Odyssey instrument (Li-COR) and ImageStudio Lite software was used to quantitate immunoblots (Li-COR).

**Immunoprecipitation**

Keratinocytes were fractionated into cytoplasmic and nuclear fractions by incubation in Buffer A [10mM HEPES, 1.5mM MgCl2, 10mM KCl, 1mM DTT] for 2 min and then diluted 1:1 in Buffer A supplemented with 0.4% NP-40 and incubated on ice for an additional 4 min. Nuclear fractions pellets were isolated by centrifugation at 4000rpm for 5 min and supernatant containing cytoplasmic material was retained for western blot analysis or discarded. The nuclear fraction was resuspended in Nuclear Lysis Buffer [50mM Tris, pH7.4, 250mM NaCl, 0.1% NP-40, 5% glycerol, protease inhibitors] and briefly sonicated at 10% power for 20 sec. Lysates were cleared by centrifugation and an aliquot was used to determine protein concentration. 100-500μg protein were diluted 1:1 in Tris buffer and 10% of lysate was retained as input. 20-60μl of M2 FLAG beads were added to lysates and incubated for 4 hours at 4°C. Beads were washed 3 times in wash buffer [50mM Tris, pH 7.4, 250mM NaCl, 0.01% NP-40, protease inhibitors] and resuspended in 2X LDS buffer (Invitrogen) after final wash. Bead were heated to 75°C for 10 min and centrifuged at max speed for 5 min, then supernatants were transferred to new tubes. Samples were run on SDS-PAGE gel for downstream western blot analysis.

**Cell Fractionation**

Cells were washed with PBS and resuspended in 500 µL of ice-cold Buffer A [10 mM HEPES, pH 7.9, 1.5 mM MgCl_2_, 10 mM KCl, 0.5 mM DTT, 1X protease inhibitor] for 2 min and then diluted 1:1 in Buffer A supplemented with 0.4% NP-40 and incubated on ice for an additional 4 min. Nuclear fractions pellets were isolated by centrifugation at 4000rpm for 5 min and supernatant containing cytoplasmic material was retained for western blot analysis. The pellet was resuspended in RIPA buffer and incubated on ice for 20 min then sonicated with a probe sonicator (Branson Digital Sonifier). Samples were centrifuged for 10 minutes at 4°C, max speed and the supernatant was saved as the nuclear fraction. Both fractions were subjected to Western blot analysis.

**Chromatin Immunoprecipitation**

Chromatin immunoprecipitation (ChIP) assays were performed as previously described with minor modifications ^10^. Briefly, human keratinocytes were cross-linked with 1% formaldehyde (Promega) and cells were lysed in nuclear lysis buffer. Chromatin was sonicated to an average fragment length of 150-250 bp using a Bioruptor (Diagenode). For ChIP-seq, a total of 240μg of chromatin was diluted in ChIP dilution buffer [0.01% SDS, 2% Triton X-100, 16.7mM Tris pH 8, 167mM NaCl], pre-cleared with Protein G Dynabead (ThermoFisher), incubated with IRF6 antibody (Novus) and chromatin was immunoprecipitated overnight at 4°C prior to purification with dynabeads. Following cross-link reversal, samples were treated with RNaseA and the DNA was purified using ChIP DNA Clean and Concentrator Kit (Zymo).

**Library Preparation**

***RNA-seq***

Libraries were prepared using the QuantSeq 3’ mRNA-Seq Library Prep Kit FWD for Illumina (Lexogen) following the manufacturer’s protocol. The quality of libraries was check with High Sensitivity DNA Bioanalyzer 2100 (Agilent) and quantification was performed using the KAPA Library Quantification Kit (Roche). Samples were run on a HiSeq 4000 instrument (Illumina) PE150 with ~15M reads per sample.

***ATAC-seq***

Cells were isolated and subjected to ATAC-seq as previously described ^11^. Briefly, 55,000 cells were pelleted after sorting and resuspended in 50μl of ATAC resuspension buffer (RSB) with 0.1% NP40, 0.1% Tween-20, and 0.01% digitonin. After three minutes, 1ml of ATAC RSB with 0.1% Tween-20 was added, tubes were inverted, and nuclei were centrifuged at 500 rcf*.* for 10 min. Supernatant was carefully removed and nuclei were resuspended in 50μl transposition mix (25ul TD buffer, 2.5μl transposase, 16.5μl PBS, 0.5μl 0.1% digitonin, 0.5μl 10% Tween-20, and 5μl water). Transposition was performed for 30 minutes at 37 C with shaking in a thermomixer at 1000 RPM. Reactions were purified with a Zymo DNA Clean & Concentrator 5 kit and library generation was performed as previously described ^11^. Samples were run on a HiSeq 4000 instrument (Illumina), PE75, ~50M reads per sample.

***ChIP-seq***

Five nanograms of DNA from chromatin immunoprecipitation was used as input for generation of libraries using the NEBNext Ultra II DNA Library Prep Kit for Illumina (NEB). Quality of DNA libraries was assessed on High Sensitivity DNA Bioanalyzer 2100 (Agilent) and library quantitation was performed using KAPA Library Quantification Kit (Roche). Samples were run on the HiSeq 4000 instrument (Illumina), PE150, ~25M reads per sample.

**Sequencing data analysis**

***General analysis of -omics data***

Reference genome hg19 and GENCODE v19 ^12^ were used. Conversions of ENSEMBL ^13^ IDs to gene symbols were done using python package biothings ^14^. Heatmaps were made using pheatmap or Treeview ^15^.

***RNA-seq analysis***

Single end reads were mapped to the hg19 reference genome with GRCh37 Ensembl annotations using STAR aligner (version 2.5.4b) ^16^ followed by generation of genes by samples counts matrices with RSEM (version 1.3.0) ^17^. Tximport ^18^ R package was using the import RSEM counts into R, and R package DESeq2 ^19^ was used to call differentially expressed genes. Finally, differential gene TPM values were visualized in heatmaps using the R package pheatmap (https://CRAN.R.project.org/package=pheatmap) or Treeview. GO term enrichment was determined via clusterProfileR ^20^ or Enrichr ^21^.

***ATAC-seq analysis***

ATAC-seq read alignment, quality filtering, duplicate removal, transposase shifting, peak calling, and signal generation were all performed through the ENCODE ATAC-seq pipeline ^22^. Briefly, adapter sequences were trimmed, sequences were mapped to the hg19 reference genome using Bowtie2 ^23^ (-X2000), poor quality reads were removed (params), PCR duplicates were removed (Picard Tools-Broad Institute, MarkDuplicates), chrM reads were removed, and read ends were shifted +4 on the positive strand or -5 on the negative strand to produce a set of filtered high-quality reads. These reads were put through MACS2 ^24^ to get peak calls and signal files. Finally, IDR analysis was run on the two replicate peak files to produce an IDR peak file that is the reproducible set of peaks across both replicates. The full pipeline can be found on the ENCODE portal. ATAC seq peaks were processed for differential expression using a pipeline described here (https://rockefelleruniversity.github.io/RU_ATAC_Workshop.html) with modifications. R package DeSeq2 ^19^ was used to determine differential ATAC peaks. ChIPseeker ^25^ was used to annotate ATAC peaks were nearest genes for GO term enrichment.

***HiChIP analysis***

Previously processed data was used for HiChIP analysis ^26^. Differential loop analysis was done using the R package diffloop ^27^.

***scRNA-seq analysis***

Previously processed data was used for scRNA-seq analysis of normal human skin (GEO 144236accession GSE) ^28^. Violin plots were generated with Seurat R package (version 3.0.0) using previously annotated normal human skin subpopulations ^28^.

***ChIP-seq analysis***

Sequencing reads were uniquely aligned to hg19 with Bowtie2 ^23^. We then filtered out reads that were unmapped, duplicate, or had less than 10 mapping quality. ChIP signals were normalized to 10 million mapped reads with peaks called using MACS2 (p-value<10^-5^, FDR<0.05, fold-enrichment>5) ^24^ on the filtered bam files, controlling ChIP samples to input samples, using the default parameters plus -f BAMPE --bdg. Peak regions across all samples were merged using bedtools ^29^. Reads in peak regions were determined using bedtools multicov, passing the bam files and the merged peak file as parameters. Further analysis was performed using the ChIPpeakAnno ^30^ R package , DESeq2 ^19^ and the annotation dataset TxDb.Hsapiens.UCSC.hg19.knownGene. For the signal density plots of ChIP-seq data, the 2 kbp around MACS2 peaks were input to deepTools computeMatrix reference-point to generate a computeMatrix file, which were then input to deepTools plotProfile and plotHeatmap. Sequences for 500 IRF6 peaks in each dataset, within ± 100bp of peak summits, were extracted for known or de novo motif analysis against these sequences using HOMER ^31^.

**Glucose transporter regulation analysis**

ATAC-seq analysis was performed as stated above to nominate transcription factors that footprint to promoter interacting regions associated with *SLC2A1, SLC2A3* and *SLC5A1*. These putative regulators were further interrogated in publicly available datasets as previously reported^32^. Briefly, regulator with knockout or knockdown data generated in keratinocytes was interrogated for expression of glucose transporters. A negative association, where gene expression decreased with regulator knockdown or knockout was reporter.

**Mass spectrometry analysis**

***LC-MS data analysis***

MS/MS data were analyzed using both Preview and Byonic v2.6.49 (ProteinMetrics). All data were first analyzed in Preview to provide recalibration criteria if necessary and then reformatted to .MGF before full analysis with Byonic. Analyses used Uniprot canonical and isoform FASTA files for Human with MT sequences concatenated as well as common contaminant proteins. Data were searched at 12 ppm mass tolerances for precursors, with 0.4 Da fragment mass tolerances assuming up to two missed cleavages and allowing for fully specific and N-ragged tryptic digestion. These data were validated at a 1% false discovery rate using typical reverse-decoy techniques ^33^. The resulting identified peptide spectral matches and assigned proteins were then exported for further analysis using custom tools developed in [MATLAB](https://www.sciencedirect.com/topics/biochemistry-genetics-and-molecular-biology/sequest) (MathWorks) to provide visualization and statistical characterization.

Spectral match assignment files were collapsed to the gene level and false positive matches and contaminants were removed. SAINT analysis ^34^ ([crapome.org](http://crapome.org)) was run with the following parameters: 10,000 iterations, LowMode ON, Normalize ON and the union of MinFold ON and OFF. Minimum interactome inclusion criteria were SAINT ≥ 0.8, fold change over control ≥ 4.

***Crosslinked peptide data analysis***

In a typical analysis, raw mass spectral data were analyzed using Byonic (Protein Metrics, San Carlos, CA, v2.14.27) to assign peptides and infer proteins. Peptide data were restricted based on tryptic digestion, allowing for n-terminal ragged cleavages, and up to two missed cleavage sites. Proteins were held to a 1% false discovery rate. Byonic X-Link functionality was used to generate predicted crosslinks between biding partners. Data were further analyzed using Byologic (Protein Metrics, San Carlos, CA) for validation, visualization, and report generation. Potential crosslinks were graded empirically based on their log probabilities, their XIC, MS1, and MS/MS spectra, as well as other qualitative features such as the presence of potential collating peptides. The crosslinked peptides were assigned a confidence ranking. High confidence crosslinks were identified as true positives or identified more than twice with some uncertainty in the peptide call (short peptide or co-isolation). For the UVC crosslinked peptide analysis, raw files were searched by Maxquant (v. 2.0.2.0) for protein identification and quantification in LFQ mode^35^ , searching against IRF6 protein sequence FASTA. The maximum number of miss-cleavages for trypsin was two per peptide. Cysteine carbamidomethylation was set as ﬁxed modification. Methionine oxidation, glucose, glucose-H_2_O and glucose-NH_3_ were set as variable modiﬁcations. The tolerances in mass accuracy were 20 ppm. The maximum false discovery rates (FDRs) were set at 0.01 at both peptide and protein levels, and the minimum required peptide length was 6 amino acids. MS/MS spectrum was plotted manually from raw files.

**QUANTIFICATION AND STATISTICAL ANALYSIS**

Statistical analyses were performed with GraphPad Prism version 9 (GraphPad Software, La Jolla, CA) or R versions 3.4.0 and 3.5.1. Parameters such as number of replicates, the number of independent experiments, measures of center, dispersion, and precision (mean ± SD or SEM), statistical test and significance, are reported in Figures and Figure Legends. Student’s T-test was used to calculate significance, unless otherwise indicated in the figure legend. For all figure panels, * indicates p-value < 0.05, ** indicates p-value < 0.01, *** indicates p-value < 0.005, and **** indicates p-value < 0.001. Metabolomics statistics were performed using a one-way ANOVA analysis and were performed by Metabolon.

**DATA AND CODE AVAILABILITY**

All raw and processed data will be made available on upon publication. Data has been deposited to GEO accession number GSE197203. The mass spectrometry proteomics data have been deposited to the ProteomeXchange Consortium via the PRIDE^36^ partner repository with the dataset identifier PXD038128 and 10.6019/PXD038128.
