## Extended Data Figures for "Glucose modulates transcription factor dimerization to enable tissue differentiation"

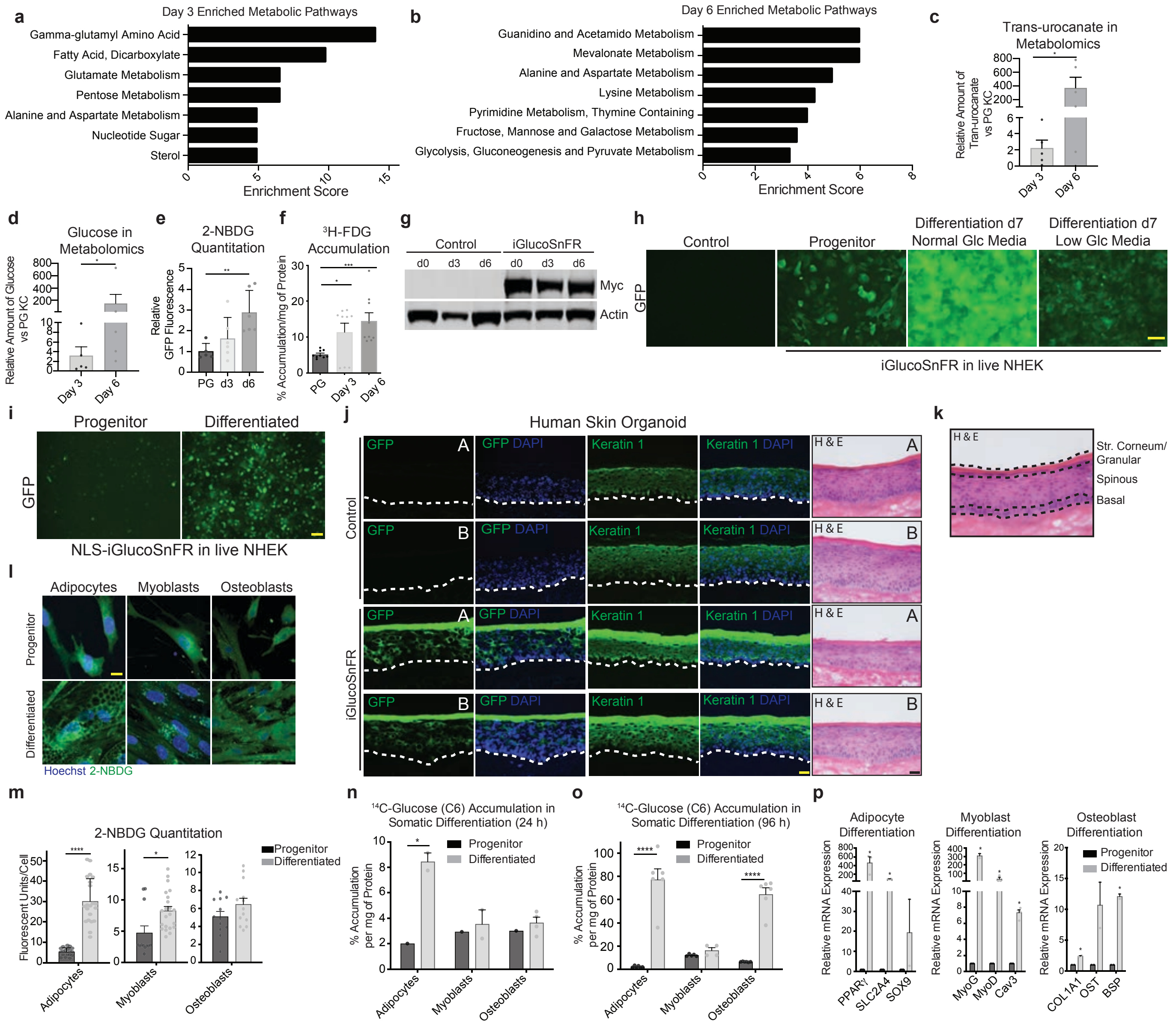


**Extended Data Fig. 1. Metabolomic analysis identifies glucose accumulation in differentiated keratinocytes.** Enriched metabolic pathways in metabolomic data from 5 mass spectrometry methods, including UPLC-MS, positive and negative ionization LC-MS, LC-polar-MS, and GC-MS, in **(a)** day 3 (d3) differentiated keratinocytes or **(b)** day 6 (d6) differentiating keratinocytes. Levels of trans-urocanate **(c)** and glucose **(d)** from MS data in differentiating keratinocytes (n=5 biological replicates). **(e)** Quantitation of 2-NBDG fluorescence in progenitor (PG), d3 or d6 differentiated keratinocytes (n=3 biological replicates, 2 fields per replicate from Fig. 1b). **(f)** Quantitation of 3H-FDG accumulation in progenitor, d3 or d6 keratinocytes (n=3 biological replicates, 3 technical replicates each). **(g)** Western blot analysis of PG, d3 or d6 keratinocytes transduced with empty vector control or the glucose sensor, iGlucoSnFR-Myc. **(h)** Representative fluorescence images of keratinocytes transduced with empty vector or glucose sensor, iGlucoSnFR-Myc in PG or d7 differentiated keratinocytes grown in standard (4.5g/L) or low (0.5g/L) glucose media. GFP signal indicates detection of free glucose (n=2 biological replicates, scale bar=20μm). **(i)** Representative fluorescence image of keratinocytes transduced with empty vector or NLS-iGlucoSnFR-Myc in PG or d7 differentiated keratinocytes. GFP signal indicates detection of free glucose in the nucleus (n=2 biological replicates, scale bar=20μm). **(j)** Immunofluorescence staining of GFP [green] and nuclear DAPI stain [blue] in epidermal organoids (d7) generated from keratinocytes transduced with empty vector control or iGlucoSnFR-Myc; hematoxylin and eosin staining of epidermal organoids is also shown (n=2 biological replicates, A and B, scale bar=20μm). **(k)** Depiction of regions delimited for the stratum corneum/granular, spinous, or basal layers for GFP quantitation of epidermal organoids in Figure 1c. Because of their adjacency and thinness, the granular layer and stratum corneum are quantified together (n=3 biological replicates, two replicates are shown, scale bar=20μm). **(l)** 2-NBDG, a fluorescent glucose analog, accumulates in adipocytes, myoblasts, and osteoblasts; 2-NBDG [green], Hoechst nuclear stain [blue] (scale bar=10μm). **(m)** Quantitation of fluorescence in (l) (n=3 biological replicates and at least 4 fields per replicate). Quantitation of radiolabeled glucose 14C-Glucose (C6) accumulation after **(n)** 24h or **(o)** 96h, normalized to mg of protein in each sample (n=3 biological replicates and 3 technical replicates per experiment). **(p)** qPCR of differentiation markers specific for adipocytes, myoblasts or osteoblasts confirming differentiation of each cell type (n=2 biological replicates). Statistical significance determined using unpaired two-tailed Student t-test. Error bars represent S.E.M., *p<0.05, **p<0.01, ***p<0.005, ****p<0.001.

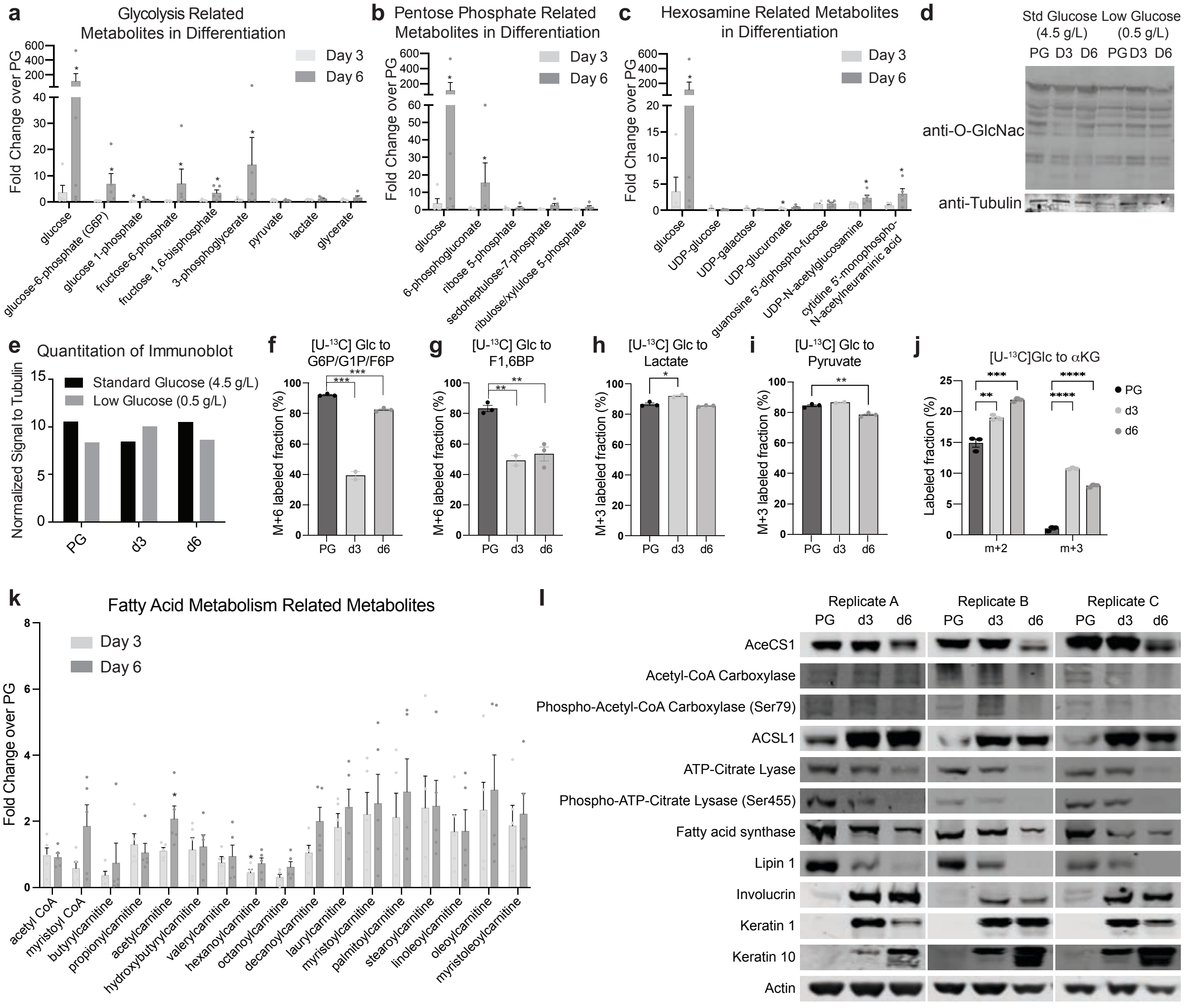


**Extended Data Figure 2. Glucose metabolism intermediates and outputs in epidermal differentiation.** Levels of metabolites related to the 3 major glucose catabolic pathways **(a)** glycolysis, **(b)** pentose phosphate, and **(c)** hexosamine from MS data in d3 or d6 differentiated keratinocytes (n=5 biological replicates). **(d)** Western blot analysis of the post-translational modification O-GlcNAc in progenitor (PG), d3 or d6 keratinocytes grown in standard (4.5g/L) or low (0.5g/L) glucose media. **(e)** Quantitation of immunoblot in (d). Metabolic flux analysis was conducted on progenitor (PG), d3 and d6 differentiated keratinocytes to evaluate [U-13C] glucose incorporation into **(f)** glucose-6-phosphate/glucose-1-phosphate/fructose-6-phosphate, **(g)** fructose-1,6-bisphosphate, **(h)** lactate, **(i)** pyruvate or **(j)** $\alpha$-ketoglutarate (n=3 biological replicates per time point). **(k)** Levels of metabolites related to fatty acid metabolism from MS data in d3 or d6 differentiated keratinocytes (n=5 biological replicates). **(l)** Western blot analysis of a panel of lipid metabolism regulators in PG, d3 and d6 differentiated keratinocytes (n=3 biological replicates). Statistical significance determined using unpaired two-tailed Student t-test. Error bars represent S.E.M., *p<0.05, **p<0.01, ***p<0.005, ****p<0.001.

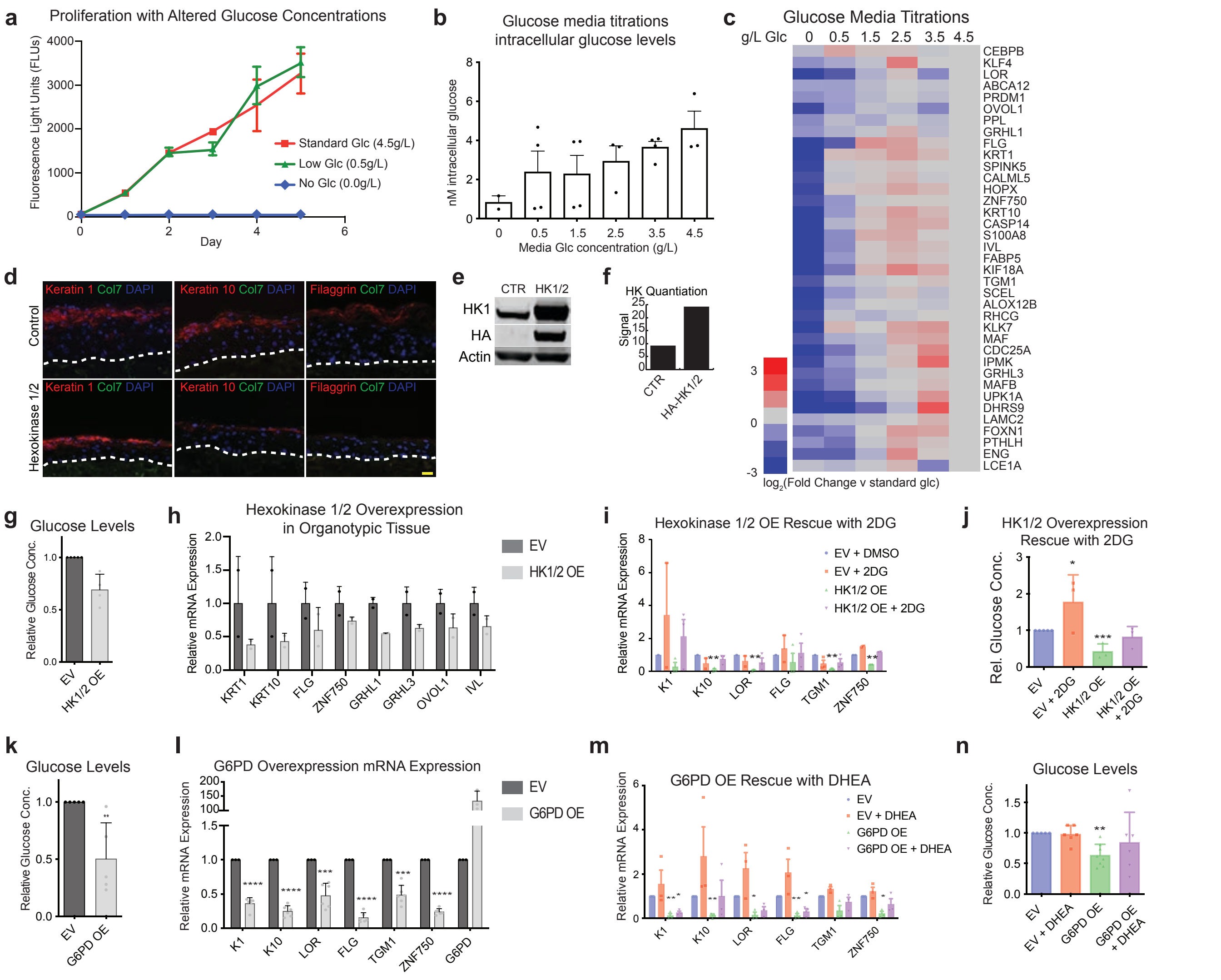


**Extended Data Figure 3. Impacts of glucose concentration on epidermal differentiation.**
**(a)** Measurement of proliferation of progenitor keratinocytes in standard(4.5g/L), low (0.5g/L) or no glucose media (n=3 biological replicates). **(b)** Intracellular glucose levels of d3 differentiated keratinocytes grown in media with varying glucose concentrations (0, 0.5, 1.5, 2.5, 3.5 or 4.5g/L) (n=3 biological replicates). **(c)** Heatmap of qPCR analysis of epidermal differentiation gene expression in varying glucose concentration media shown in (b). **(d)** Immunostaining of differentiation markers [red], collagen VII [green], and nuclear DAPI stain [blue] in epidermal organoids generated with keratinocytes transduced with empty vector (EV) control or HA-hexokinase 1 and 2 (HK1/2) (n=2 biological replicates, scale bar=20$\mu$m). **(e)** Western blot analysis of keratinocytes transduced with EV control or HK1/2, along with quantitation of immunoblot in **(f)**. **(g)** Intracellular glucose levels in differentiated keratinocytes overexpressing HK1/2 (n=2 biological replicates). **(h)** qPCR of epidermal differentiation gene expression in epidermal organoids overexpressing HK1/2 (n=2 biological replicates). **(i)** qPCR of epidermal differentiation gene expression in differentiated keratinocytes (d3) transduced with EV control or HK1/2 treated with DMSO or 2-deoxy-d-glucose (2DG) (n=2 biological replicates). **(j)** Measurement of intracellular glucose levels in (i) (n=2 biological replicates). **(k)** Intracellular glucose levels in differentiated keratinocytes transduced with empty vector control or G6PD (n=2 biological replicates). **(l)** qPCR of epidermal differentiation gene expression in d3 differentiated keratinocytes transduced with EV control or G6PD (n=2 biological replicates). **(m)** qPCR of epidermal differentiation gene expression in d3 differentiated keratinocytes transduced with EV control or G6PD treated with DMSO or the DHEA inhibitor of G6PD (n=2 biological replicates). **(n)** Measurement of intracellular glucose levels in (m) (n=2 biological replicates). Statistical significance determined using unpaired two-tailed Student t-test. Error bars represent S.D., *p<0.05, **p<0.01, ***p<0.005, ****p<0.001.

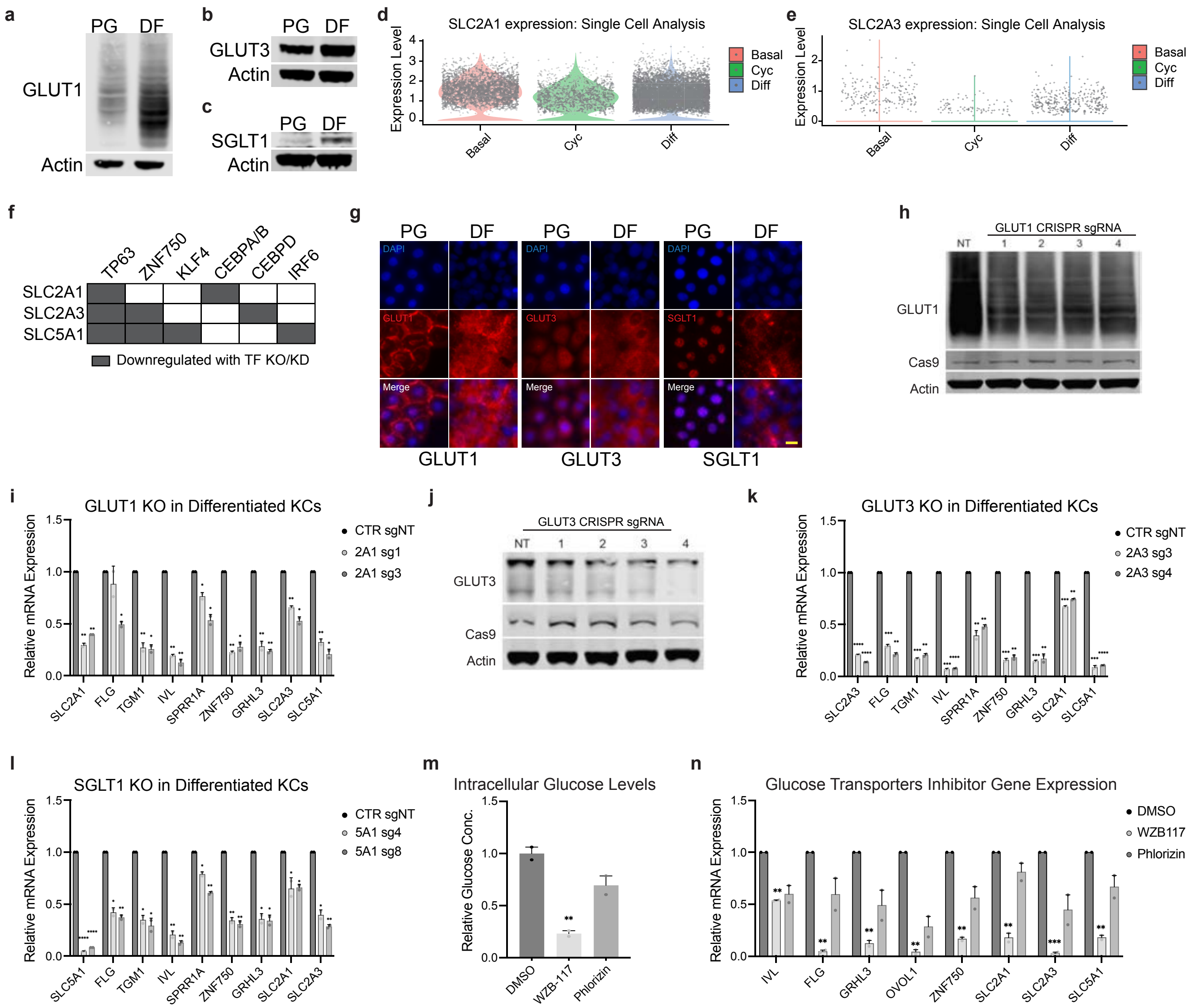


**Extended Data Figure 4. Glucose transporters in epidermal differentiation.**

Western blot analysis of **(a)** GLUT1, **(b)** GLUT3 or **(c)** SGLT1 in progenitor (PG) and d3 differentiated keratinocytes (GLUT1 protein runs as a broad band on western analysis). Expression of **(d)** GLUT1 (*SLC2A1*) and **(e)** GLUT3 (*SLC2A3*) in single-cell RNA-seq data from normal human epidermis in the basal, cycling and differentiated cell populations. **(f)** Analysis of *SLC2A1*, *SLC2A3*, and *SLC5A1* regulation by epidermal transcription factors (TFs); gray represents gene expression downregulation with TF knockdown or knockout in keratinocytes interrogated in publicly available data compiled in Lopez-Pajares et al. **(g)** Immunofluorescence staining of glucose transporters [red] and nuclear DAPI stain [blue] in PG and d3 differentiated keratinocytes (scale bar=10μm). **(h)** Western blot analysis of GLUT1 protein levels in keratinocytes treated with non-targeting (NT) or sgRNAs targeting *SLC2A1* (GLUT1). **(i)** qPCR of differentiation gene expression in *SLC2A1* KO d3 differentiated keratinocytes (n=2 biological replicates). **(j)** Western blot analysis of GLUT3 protein levels in keratinocytes treated with NT or sgRNAs targeting *SLC2A3* (GLUT3). **(k)** qPCR of differentiation gene expression in *SLC2A3* KO d3 differentiated keratinocytes (n=2 biological replicates). **(l)** qPCR of differentiation gene expression in *SLC5A1* (SGLT1) KO keratinocytes (n=2 biological replicates). **(m)** Relative intracellular glucose concentrations in differentiated keratinocytes treated with DMSO, GLUT inhibitor WZB117 (10μM), or SGLT inhibitor phlorizin (50μM) (n=3 biological replicates). **(n)** qPCR analysis of differentiation gene expression in differentiated keratinocytes treated with DMSO, WZB117 or phlorizin (n=3 biological replicates). Statistical significance determined using unpaired two-tailed Student t-test. Error bars represent S.E.M., *p<0.05, **p<0.01, ***p<0.005.

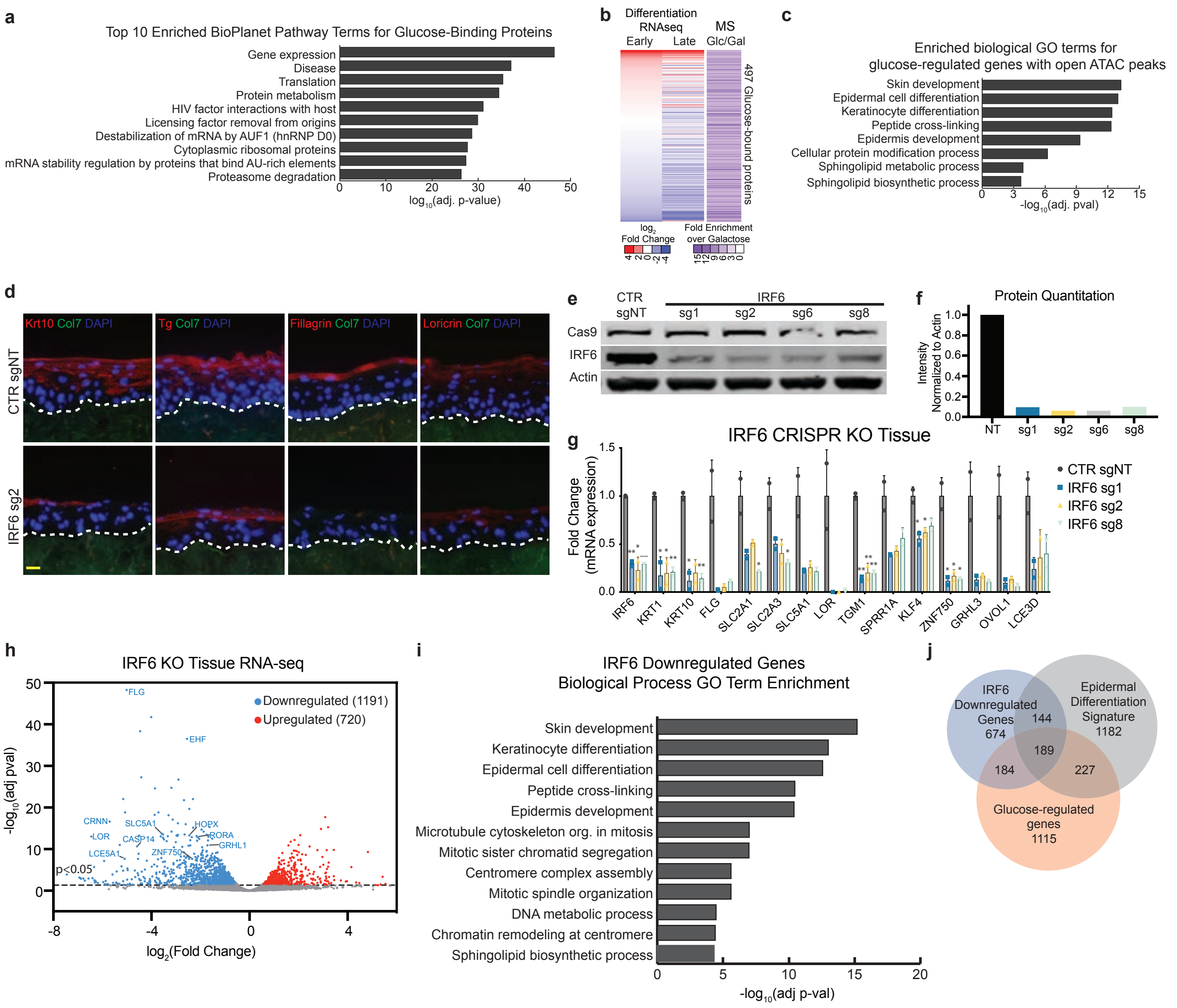


**Extended Data Figure 5. Identification of IRF6 as a glucose-binding pro-differentiation transcription factor.**

**(a)** Top 10 terms for pathway enrichment using the BioPlanet database for 497 glucose-binding proteins (Enrichr analysis). **(b)** Heatmap of 497 proteins identified as glucose-binding proteins showing mRNA expression from RNA-seq in early (d3) and late (d6) differentiated keratinocytes, and enrichment in glucose elution versus galactose elution in dextran column affinity chromatography followed by LC-MS/MS. **(c)** Enriched GO terms for glucose-regulated genes with open (increased) ATAC-seq peaks. **(d)** Epidermal organoids of primary human keratinocytes transduced with Cas9 and non-targeting (NT) control sgRNA or *IRF6*-targeting sgRNA representing *IRF6* knock-out (KO) human tissue harvested at d7; differentiation markers [red], nuclei [blue], collagen VII basement membrane staining [green] (n=3 biological replicates, representative image shown, scale bar=20μm). **(e)** Western blot analysis of keratinocytes transduced with Cas9 and control NT sgRNA or 4 different *IRF6* targeting sgRNAs, used to generate epidermal organoids. **(f)** Quantitation of IRF6 protein levels in (e). **(g)** Differentiation gene expression in *IRF6* KO tissue measured by qPCR (n=2 biological replicates per sample). **(h)** Volcano plot of PAS-seq data from *IRF6* KO tissue showing 720 upregulated and 1191 downregulated genes (n=3 biological replicates, FDR>0.05, fold change>1.5). Highlighted are downregulated differentiation-specific genes. **(i)** Enriched biological process GO terms for *IRF6* KO tissue downregulated genes. **(j)** Overlap of *IRF6* KO downregulated genes with glucose-regulated genes and the epidermal differentiation gene signatures generated by Lopez-Pajares et al. and Rubin et al. Statistical significance determined using unpaired two-tailed Student t-test. Error bars represent S.D., *p<0.05, **p<0.01, ***p<0.005, ****p<0.001.

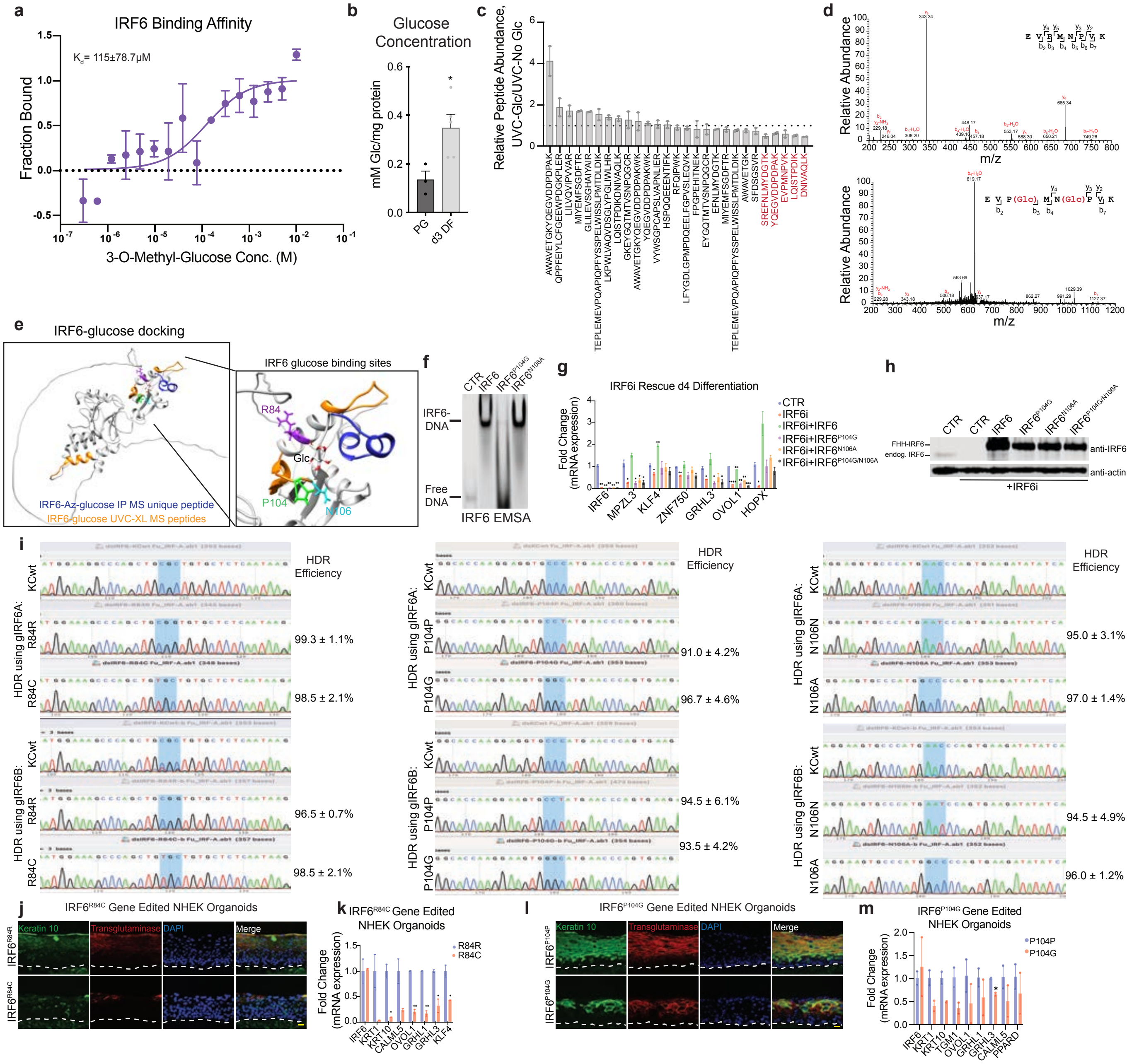

**Extended Data Figure 6. IRF6 glucose binding capacity is required for epidermal differentiation.** **(a)** Microscale thermophoresis (MST) analysis of IRF6 affinity toward the metabolically stable glucose analog, 3-O-methyl-glucose (Kd=115±78.7μM) (n=3). **(b)** Quantitation of intracellular glucose levels in progenitor and d3 differentiated keratinocytes; (n=5 biological replicates). **(c)** Candidate IRF6 peptides enriched for glucose binding (n=2). **(d)** MS/MS of the IRF6 peptide identifying the P104 and N106 residue crosslinked to glucose. **(e)** Modeling the IRF6-glucose interaction using Autodock Vina, showing a highly scored model (score<-5) and the location of the unique peptide identified via azido-glucose IP-MS (blue) and peptides identified via UV-C crosslinking of glucose to IRF6 followed by MS (orange); the enlarged image shows the location of disease-associated residue arginine 84 (purple), as well as proline 104 (green) and asparagine 106 (cyan) identified as glucose-binding residues. **(f)** Electrophoretic mobility shift assay of recombinant IRF6 and P104G or N106A glucose-binding deficient mutants. **(g)** Gene expression of d4 differentiated keratinocytes treated with control or IRF6 siRNA with enforced expression of Flag-HA-6XHis (FHH) tagged-WT, P104G, N106A or P104G/N106A IRF6 constructs to rescue the IRF6i differentiation defect (representative technical replicates shown, n=2 biological replicates). **(h)** Western blot analysis of samples in (g). **(i)** Sanger sequencing chromatogram of targeted editing sequence at the endogenous IRF6 gene locus; editing efficiencies are noted for each of the two replicates (n=2 biological replicates). **(j)** Immunostaining of keratin 10 [green], transglutaminase [red], and nuclear DAPI [blue] in epidermal organoids generated from R84R and R84C gene edited keratinocytes; (shown is a representative image from 2 biological replicates, scale bar=20μm). **(k)** Gene expression analysis of R84R and R84C gene edited tissue (n=2 biological replicates). **(l)** Immunostaining of keratin 10 [green], transglutaminase [red], and nuclear DAPI [blue] in epidermal organoids generated from P104P and P104G gene edited keratinocytes (shown is a representative image from 2 biological replicates, scale bar=20μm). **(m)** Gene expression analysis of P104P and P104G gene edited tissue (n=2 biological replicates). Statistical significance determined using unpaired two-tailed Student t-test. Error bars represent S.E.M., *p<0.05, **p<0.01, ***p<0.005, ****p<0.001.

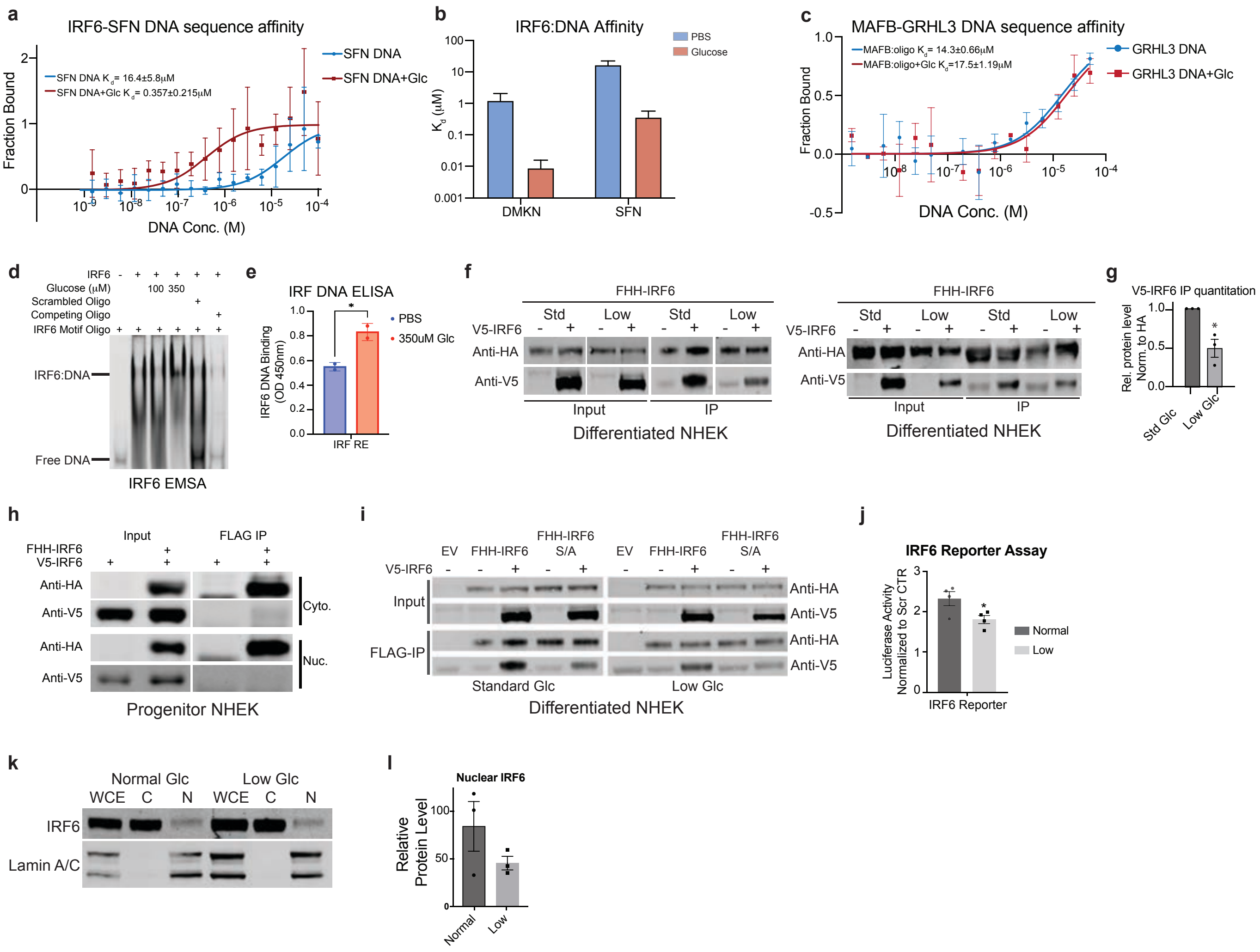


### Extended Data Figure 7. Functional impacts of glucose on IRF6.

**(a)** Affinity of IRF6 towards cognate DNA binding sequence in the SFN locus measured by MST. Measurements were conducted using a wild-type SFN sequence DNA oligo in PBS (Kd=16.4±5.8μM) or 350μM glucose (Kd=0.357±0.215μM) (n=4). **(b)** Summary of the change in affinity of IRF6 toward DNA ± 350μM glucose, log scale shown. **(c)** Affinity of MAFB towards cognate DNA binding sequence in the GRHL3 locus measured by MST. Measurements were conducted using a wild-type GRHL3 sequence oligo in PBS (Kd=14.3±0.66μM) or 350μM glucose (Kd=17.5±1.19μM) (n=3). **(d)** Electrophoretic mobility shift assay of IRF6 motif probe incubated with recombinant IRF6 with or without glucose or incubation with scrambled or competing oligonucleotides. Indicated are the free DNA and IRF6-DNA complex. **(e)** DNA ELISA of recombinant IRF6 binding to an IRF responsive element with or without glucose (n=2). **(f)** Western blot analysis of replicate samples of FHH-IRF6 co-immunoprecipitation from FHH-IRF6 and V5-IRF6 transduced keratinocytes differentiated for 3 days in standard(4.5g/L) or low (0.5g/L) glucose media and quantitation **(g)** (n=2 biological replicates). **(h)** Western blot analysis of FHH-IRF6 co-immunoprecipitation from FHH-IRF6 and V5-IRF6 transduced progenitor keratinocytes in cytoplasmic or nuclear cellular fractions. **(i)** Western blot analysis of FHH-IRF6 or FHH-IRF6S413A/S424A mutant (S/A) and V5-IRF6 transduced keratinocytes differentiated in standard or low glucose media for 3 days after FLAG immunoprecipitation. **(j)** IRF6 luciferase reporter assay for d3 differentiated keratinocytes grown in standard or low glucose media (n=4 biological replicates). **(k)** Western blot analysis of cellular fractionation of d3 differentiated keratinocytes grown in standard or low glucose media along with **(l)** quantitation of IRF6 protein levels in nuclear extracts. Statistical significance determined using unpaired two-tailed Student t-test. Error bars represent S.E.M., *p<0.05.

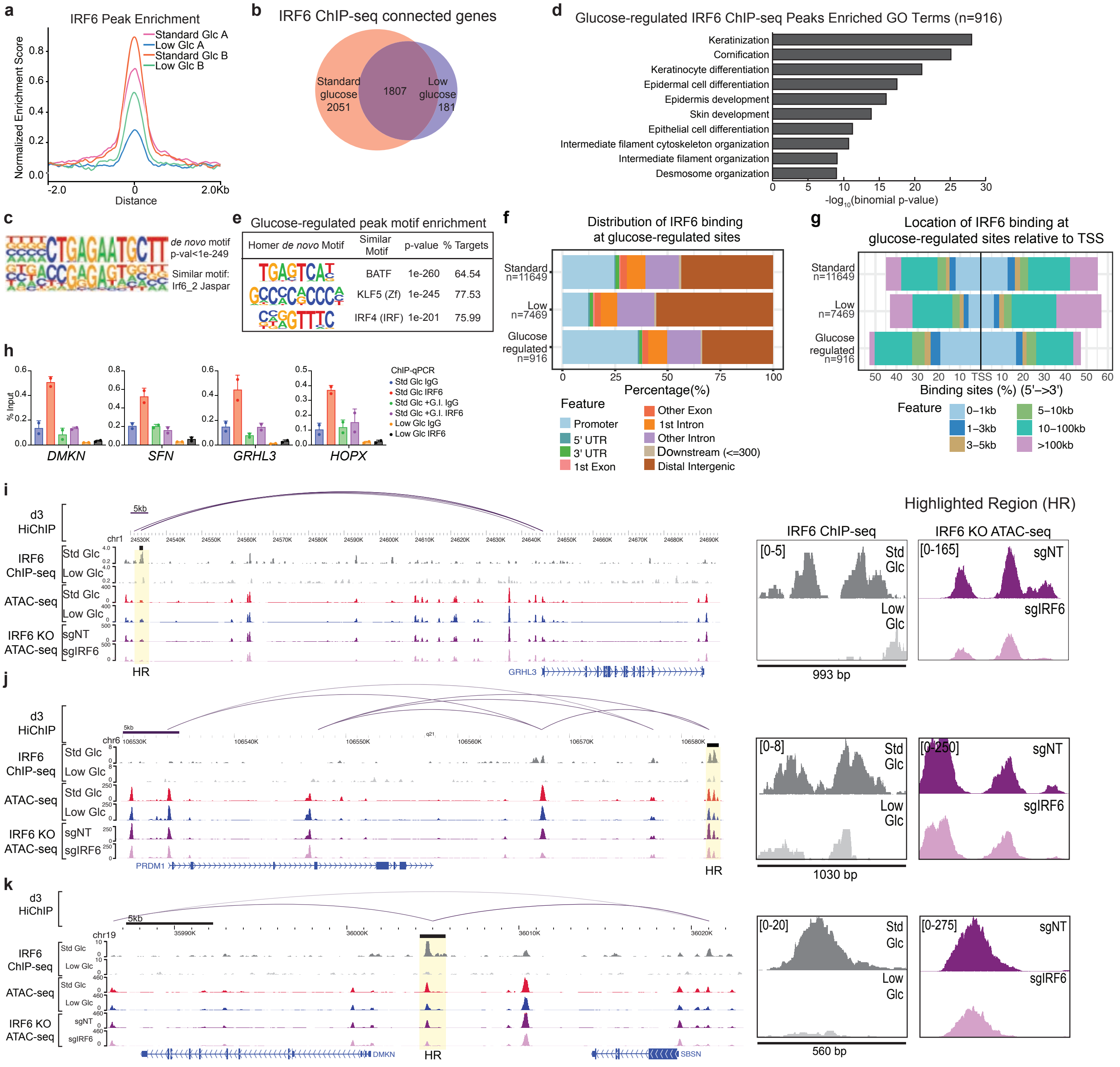


### Extended Data Figure 8. Glucose-mediated IRF6 gene regulation.

**(a)** Normalized enrichment score for IRF6 ChIP-seq peaks in differentiated keratinocytes grown in standard or low glucose media (n=2 biological replicates). **(b)** Overlap of genes connected (cGenes) to IRF6 ChIP-seq peaks in standard (3858 genes) or low (1988 genes) glucose media (n=2 biological replicates). **(c)** HOMER de novo motif analysis of shared IRF6 ChIP-seq peaks in standard and low glucose media compared to known IRF6 Ѧ SPAR motif 2. **(d)** Gene ontology (GO) terms for glucose-modulated IRF6 ChIP-seq peaks identified in standard or low glucose conditions. **(e)** Top three enriched HOMER de novo motifs for 916 glucose-regulated ChIP-seq peaks. **(f)** Genomic features associated with standard, low or glucose-regulated IRF6 ChIP-seq peaks. **(g)** Location of IRF6 binding relative to TSS in standard, low or glucose-regulated ChIP-seq peaks. **(h)** IRF6 chromatin immunoprecipitation followed by qPCR of the *DMKN*, *SFN*, *GRHL3* and *HOPX* loci after d3 differentiation in standard glucose media (4.5g/L), standard glucose media with GLUT and SGLT1 inhibitors (10μM WZB-117 and 50μM phlorizin) treated for 15 h, or low glucose media (0.5g/L) (representative data shown is 2 technical replicates, n=2 biological replicates). Genome browser tracks of IRF6 ChIP-seq peaks and ATAC-seq peaks in d3 differentiated keratinocytes in standard or low glucose media, and ATAC-seq peaks in control (NT) or *IRF6* KO keratinocytes for the **(i)** *GRHL3*, **(j)** *PRDM1*, and **(k)** *SBSN/DMKN* loci. Error bars represent S.E.M.

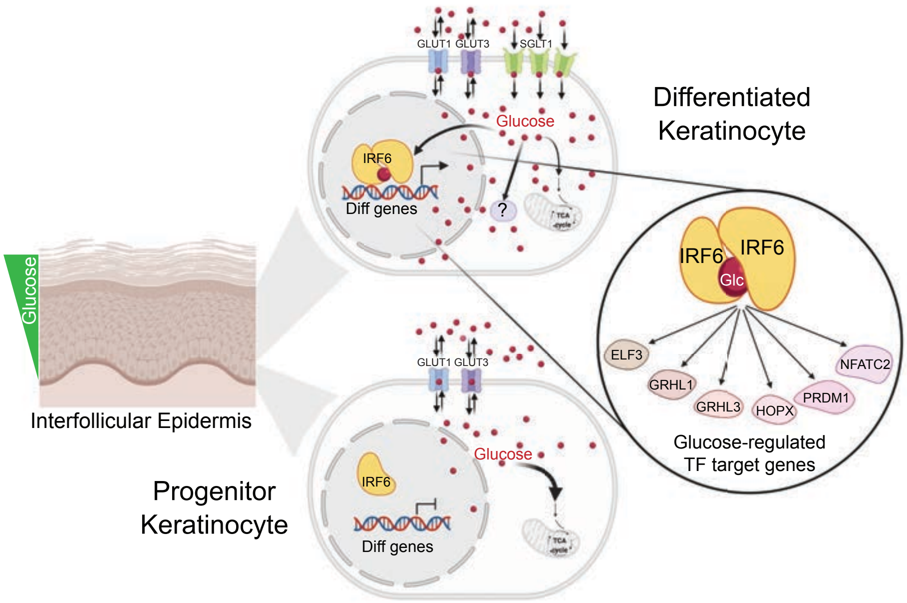


**Extended Data Figure 9. Model of glucose-mediated gene regulation in epidermal differentiation.** Glucose accumulation is a feature of epidermal differentiation and free glucose accumulates in the differentiated layers of the epidermis. In progenitor keratinocytes, glucose equilibrium is maintained by the facilitative glucose transporters, GLUT1 and GLUT3 and glucose is metabolized. Upon differentiation, the sodium-coupled glucose co-transporter, SGLT1, is induced and glucose accumulation occurs, without increased glucose catabolism. Glucose binds the IRF6 pro-differentiation TF. Glucose enhances IRF6 dimerization and genomic targeting to regulate gene expression of IRF6-dependent target genes, including pro-differentiation transcription factors.
